# *Toxoplasma gondii* GRA66 prevents premature egress driven by the host phospholipase RARRES3, independently of RNF213

**DOI:** 10.64898/2026.09.12.751112

**Authors:** Caroline de Moraes de Siqueira, Stephanie Martinez-Beltran, Emma A Johnston, Jörn Coers, Jeroen P. J. Saeij

## Abstract

*Toxoplasma gondii* replicates inside a host-derived parasitophorous vacuole (PV), and interferon-gamma (IFNγ) induces host restriction factors that target this compartment. Previous CRISPR screens identified the dense granule protein GRA66 and the GRA57/GRA70/GRA71 complex as required for parasite fitness in IFNγ-stimulated human cells, but the host pathways they oppose were unknown. Here, we show that GRA66, a PV membrane (PVM)-associated protein predicted to be an N-acylphosphatidylethanolamine (NAPE)-hydrolyzing phospholipase D, is required to prevent premature egress driven by the host phospholipase and acyltransferase RARRES3. Loss of GRA66 caused premature parasite egress, host cell death, and impaired replication in human cells. These phenotypes persisted in cells lacking RNF213, the E3 ubiquitin ligase that dominates IFNγ-dependent *Toxoplasma* restriction in human cells, and *Δgra66* vacuoles recruited less RNF213 and ubiquitin than wild type, indicating an RNF213-independent mechanism rather than an exaggerated RNF213 response. Complementation with a catalytic-site mutant of the GRA66 zinc-binding motif failed to restore any of these phenotypes, indicating a requirement for its predicted enzymatic activity. Deleting RARRES3 rescued the premature egress of Δ*gra66* and catalytic-mutant parasites but not of *Δgra70* parasites, whereas the replication defect persisted, revealing a second, RARRES3-independent consequence of GRA66 loss. RARRES3 was recruited to the PVM and intravacuolar network after IFNγ stimulation, and structural modeling supported assignment of GRA66 to the NAPE-phospholipase D family with an intact di-zinc active site. These findings define a lipid-centered, RNF213-independent arm of human cell-autonomous immunity and identify the parasite effector required to withstand it.

**Importance:** *Toxoplasma gondii* infects a large fraction of the world’s population and persists for life inside host cells, sheltered within a compartment called a vacuole. The immune signal interferon gamma normally holds the parasite in check, but how human cells attack this vacuole, and how the parasite resists, is only partly understood because humans lack the main anti-vacuolar defenses mice use. Here, we show that human cells recruit a lipid-remodeling enzyme, RARRES3, to this vacuole, and that RARRES3 drives the parasite to exit prematurely, triggering host cell death. The parasite counters with a lipid-modifying enzyme of its own, GRA66, which is required to prevent that premature exit. This host defense works independently of the pathway thought to dominate human control of *Toxoplasma*. Our findings identify lipid remodeling of the pathogen-containing vacuole as a distinct arm of human cell-intrinsic immunity, and the parasite enzyme that counteracts it as a potential drug target.

## Introduction

Pathogen-containing vacuoles provide a specialized intracellular niche that allows pathogens to acquire nutrients while limiting immune detection (1, 2). To establish and maintain this niche, pathogens deploy effector proteins that remodel vacuole composition and trafficking, thereby preventing lysosomal degradation and immune recognition (3). Whereas bacteria such as *Mycobacterium* and *Salmonella* block vacuole maturation at different stages of the endocytic pathway (1), apicomplexan parasites such as *Toxoplasma gondii* and *Plasmodium spp*. create non-fusogenic vacuoles by secreting their own specialized proteins, excluding host endocytic machinery (2, 4–8). Regardless of the specific strategy employed, remodeling the vacuole is essential for intracellular survival and replication.

Following invasion, *Toxoplasma* remodels the parasitophorous vacuole (PV) through the sequential secretion and incorporation of rhoptry proteins (ROPs) and dense granule proteins (GRAs), which shape the vacuole and its interactions with the host (9, 10). GRAs localize to the PV membrane (PVM), the vacuolar lumen, or are exported into the host cell, where they perform distinct functions. At the PVM, GRA17, GRA23, GRA47, and GRA72 influence small molecule permeability (11, 12), whereas exported GRAs such as *Tg*IST and *Tg*NSM suppress IFNγ-induced host responses (13, 14). Correct delivery of GRAs to the PVM is itself an active process, as GRA45 has a chaperone-like domain required for proper GRA localization to the PVM (15). Together, these studies demonstrate that GRAs preserve the intracellular niche by maintaining PV function and limiting host immunity.

In previous genome-wide CRISPR-Cas9 screens, GRA66 and members of a heterotrimeric complex comprising GRA57, GRA70, and GRA71 were identified as parasite effectors required for fitness in IFNγ-stimulated human cells (14, 15); genome-wide screens in IFNγ-activated murine macrophages have likewise identified GRA effectors required for parasite fitness (15). GRA66 is a putative N-acyl phosphatidylethanolamine phospholipase D (NAPE-PLD) that localizes to the PV lumen and PVM (16, 18). Loss of GRA66 increases parasite susceptibility to IFNγ and triggers premature parasite egress in human foreskin fibroblasts (HFFs) (16). Members of the GRA70 complex (GRA70, GRA57, and GRA71) similarly localize to the PV and PVM and are required for optimal parasite growth during IFNγ stimulation, with loss of any member producing a comparable premature egress phenotype (16, 17). These findings suggested that GRA66 and the GRA70 complex protect intracellular parasites from IFNγ-induced restriction, but the host pathways they target remain unknown.

IFNγ induces hundreds of interferon-stimulated genes (ISGs) that mediate cell-autonomous immunity (19, 20), yet the mechanisms that restrict *Toxoplasma* in human non-immune cells remain incompletely understood (21). In murine cells, *Toxoplasma* restriction is mediated primarily by immunity-related GTPases (IRGs) and guanylate-binding proteins (GBPs), which accumulate on the PV and promote membrane disruption and parasite elimination (22–25). Humans lack the expanded IRG repertoire found in mice (26), and although human GBPs contribute to cell-autonomous pathogen control, they are not detectably recruited to the PV in IFNγ-stimulated lung carcinoma epithelial A549 cells (26), suggesting that distinct ISG-dependent mechanisms mediate parasite restriction at the PVM-cytosol interface in human cells.

Recent studies identified Ring Finger Protein 213 (RNF213) as a key mediator of IFNγ-induced *Toxoplasma* restriction in human fibroblasts (HFFs), the human monocytic cell line THP-1, and epithelial A549 cells (27–29). RNF213 is a large E3 ubiquitin ligase that is recruited to the PV, where it promotes ubiquitin deposition and the subsequent recruitment of ubiquitin-binding adaptors, including p62 and NDP52, thereby restricting parasite replication (27–29). Consistent with this, ubiquitination of the vacuole and recruitment of p62 and NDP52 restrict parasite growth in IFNγ-activated human cells in a strain-dependent manner (30).

Whereas RNF213 defines a ubiquitin-dependent arm of this response, an independent overexpression screen for IFNγ-induced human restriction factors performed in A549 and HeLa cells identified the phospholipase and acyltransferase Retinoic Acid Receptor Responder 3 (RARRES3/PLAAT4), as a IFNγ-induced inhibitor of *Toxoplasma* growth (31). RARRES3 localizes to cellular membranes, where it remodels phospholipids through its phospholipase and acyltransferase activities (32, 33). Beyond its established roles in retinoic acid signaling and tumor suppression (34), RARRES3 has emerged as an important ISG with antimicrobial activity against both viral pathogens and *Toxoplasma* (31, 35). RARRES3 promotes premature parasite egress through its enzymatic activity whereas the catalytic mutant C113A no longer restricts the parasite, but the mechanism by which it restricts intracellular parasites remains unknown. These observations raised the possibility that parasite effectors required for fitness in IFNγ-stimulated cells counteract RNF213- or RARRES3-mediated restriction.

Here, we identify GRA66 as a parasite effector that prevents RARRES3-dependent premature parasite egress through a mechanism that is independent of RNF213. We show that an intact predicted catalytic motif of GRA66 is required to prevent premature parasite egress, and that structural modeling supports its assignment as a NAPE-PLD with an intact di-zinc active site. Furthermore, we demonstrate that RARRES3 is recruited to the PVM following IFNγ stimulation, identifying a previously unrecognized host pathway that targets the *Toxoplasma* vacuole and defining a lipid-centered, RNF213-independent arm of human cell-autonomous immunity.

## Methods

### Host Cells and Parasite Culture

*Toxoplasma* strains (Table S1) were maintained in monolayers of HFFs. HFFs and lung carcinoma epithelial (A549) cells were cultured using Dulbecco’s Modified Eagle Medium (DMEM) supplemented with 10% fetal bovine serum (FBS), 2 mM L-Glutamine, 100 μg/mL streptomycin, 100 U/mL penicillin, and 10 μg/mL gentamicin. Parasites were maintained in a similarly prepared DMEM medium supplemented with 5% FBS and no gentamicin. Cultures were incubated at 37°C with 5% CO2 as previously described (36).

### Immunofluorescence Assay: PVM GRA Localization

For dense granule protein localization, HFFs or A549 cells were seeded on 12mm glass coverslips in 24-well plates and grown to ∼90% confluency before infection with syringe-lysed wild-type, knockout, or complemented strains at an MOI of 1 for 24h. For GRA23, parasites were transiently transfected with the pGRA1-GRA23-HA-FLAG plasmid (12). After 24h, the coverslips were washed 3 times with 1X phosphate-buffered saline (PBS) to remove extracellular parasites and fixed with a solution of 4% paraformaldehyde (PFA) in PBS for 20 minutes or 4% PFA/0.001% glutaraldehyde for 1h followed by quenching with 0.1 M glycine, only for GRA66 localization. The cells were permeabilized with a 0.02% saponin or 0.2% Triton buffer made with PBS for 20 minutes and blocked with a 3%FBS/4%BSA buffer in PBS. The blocked cells were incubated overnight at 4°C with primary antibodies: rat anti-HA (1:500; Sigma-Aldrich, St. Louis, MO, USA, cat. no. 1186743100), rabbit anti-GRA23 (kindly provided by Dr. Tatsunori Masatani, Gifu University, Japan), rabbit anti-DYKDDDDK Tag (Cell Signaling, Danvers, MA, USA, cat. no. 2368S), or mouse anti-GRA7 (1:1000-1:5000; kindly provided by John Boothroyd’s Lab). After this overnight incubation, the cells were stained for 1 h at room temperature with the following secondary antibodies (1:3000): goat anti-rat 594 (Alexa Fluor, Thermo Fisher Scientific, Waltham, MA, USA, cat. no. A-11007), goat anti-mouse 594 (Alexa Fluor, Thermo Fisher Scientific, cat. no. A-11005), or donkey anti-mouse 488 (Alexa Fluor, Thermo Fisher Scientific, cat. no. A-21202). Nuclei were stained using DAPI (Thermo Fisher Scientific, cat. no. D1306). The slides were prepared using a Mowiol mounting medium prepared with Mowiol 4-88, Tris-Cl (0.2 M, pH 8.5), 1,4-diazabicyclo-[2,2,2]-octane (DABCO) in glycerol. The slides were subsequently analyzed with the NIS-Elements software (Nikon, Tokyo, Japan) and a CoolSNAP EZ digital camera (Roper Scientific, Tucson, AZ, USA), connected to an inverted fluorescence microscope (Eclipse Ti-S; Nikon, Tokyo, Japan) or Leica TCS SP8 STED 3X (Leica Microsystems, Wetzlar, Germany) using Leica LAS X software. For GRA23 localization, the slides were prepared in a blinded manner. To prevent observer bias, all samples were randomly coded with numerical identifiers.The key to the numerical codes was only unveiled after all data collection was finalized. At least 50 vacuoles were counted. Vacuoles were categorized into two groups based on GRA23 localization: (i) PVM localization, defined by a ring around the vacuole with a complete absence of signal within the PV lumen; and (ii) PV lumen localization, characterized by either exclusive luminal signal or a predominantly stronger signal in the lumen compared to the PVM.

### Generation of Parasite Strains

For generation of GRA66 complemented parasites and endogenous tagging by transfection, parasites were manually syringe lysed out of HFF monolayers grown in T25 cell culture flasks by passing the cell solution twice through 27G needles. The parasites were washed with cytomix buffer (2 mM EDTA, 120 mM KCl, 0.15 mM CaCl_2_, 10 mM K_2_HPO_4_/KH_2_PO_4_, 25 mM HEPES, and 5 mM MgCl_2_, pH 7.4), and 1x10^7^ parasites were transfected with the appropriate plasmid and/or template for gene editing, using Gene Pulser Xcell™ (Bio-Rad Laboratories, Hercules, CA, USA) as previously described (37). For GRA66 complementation, the 5’ UTR, 3’UTR and full CDS of *gra66* gene were amplified from the genomic DNA of RHΔ*hxgprt* LUC+ parasites using the primers described in Table S2 adding a C-terminal HA tag, and cloned into pUPRT::DHFR-D (Addgene, cat. no. 58528) plasmid by Gibson Assembly^®^ (New England Biolabs, cat. no.E5510S, Ipswich, MA, USA), according to manufacturer’s instructions, generating the plasmid pUPRT::GRA66-HA. This plasmid was used for generating point mutations in the predicted active site using Q5^®^ Site-Directed Mutagenesis Kit (New England Biolabs, Cat #E0554S, Ipswich, MA, USA) following the manufacturer’s instructions. Point mutations were made by replacing Histidine 540 (CAC) by Alanine (GCT) and Histidine 542 (CAC) by Alanine (GCC). The resulting plasmid (pUPRT::GRA66mut-HA) was confirmed by Sanger sequencing. Plasmids for complementation were transfected in *Δgra66* parasites (16). For GRA66 endogenous tagging, 1014 bp of the *GRA66* gene was amplified by PCR, excluding the stop codon and cloned into pLIC_3xHA_DHFR (38), adding a C-terminal 3xHA tag. The guide RNA was designed using EuPaGDT (Eukaryotic Pathogen gRNA Design Tool) (39) and cloned into pss013 (40), generating the plasmid pss013sgRNA#118. The pLIC_GRA66-3xHA plasmid was linearized with NsiI enzyme (New England Biolabs, cat. no.R3127S) and co-transfected with pss013sgRNA#118 in RH*Δku80Δhpt* parasites. The transfected parasites were selected with 10 μM FUDR (5-fluorodeoxyuridine), for complementation in the *UPRT* locus, or 2 μM pyrimethamine (for DHFR containing plasmids) and cloned by limiting dilution. All plasmids used in this study were described in Table S4.

### Production of Lentivirus and Generation of A549 Cell Lines

A549 ΔRNF213 cells were from Dr. Jörn Coers (28) and ΔRARRES3 were from Dr. David Sibley (31), as well as the plasmid with the sgRNA to generate the ΔRARRES3 line. For lentivirus production, Lenti-X 293T cells (kindly provided by Dr. Bennet Penn, University of California Davis) were grown to 70% confluency in 6-well plates. Afterwards, 1 μg of pLentiCRISPR-RARRES3 was prepared along with 250 ng pMD2G, 250 ng pRSV-Rev, and 500 ng pMDLg/RRE (gift from Dr. David Sibley, Washington University in St. Louis) (31), which were mixed with the PolyJet™ in vitro DNA transfection reagent (SignaGen Laboratories, Frederick, MD, USA) following the manufacturer’s instructions. The supernatant containing the lentiviruses was harvested at 48h and 72h and combined with 4 μg/mL Polybrene (Sigma-Aldrich, St. Louis, MO, USA) and 50 mM HEPES and spun down at 800 × g for 5 minutes. Afterwards, the supernatant was collected and used immediately or stored at -80 °C. To generate the double knockout A549 ΔRNF213/ΔRARRES3 cells, the pLentiCRISPR-RARRES3 was transduced into ΔRNF213 cells. 7 × 10^4^ A549 ΔRNF213 cells were seeded into 24-well plates. The next day, the regular media was changed to a medium containing 4 μg/mL of Polybrene, 3% FBS, and 25 mM HEPES. 500 μL of the lentivirus supernatant was added to the cells and spinoculated at 800 × g for 45 minutes at 37 °C. 6h after the spinoculation, the medium was replaced with DMEM with 10% FBS, 2 mM L-glutamine, 100 μg/mL streptomycin, 100 U/ml penicillin, and 0.1 mM of nonessential amino acids. 48h later, the cells were split and selected for at least two weeks with 1 μg/mL of Puromycin before limiting dilution cloning. Successful knockout clones were validated by PCR amplification of the target locus and subsequent confirmation by Sanger sequencing, where deletions were found in both alleles. For RARRES3 overexpression and complementation TRIP.RARRES3 or TRIP.GFP plasmid (31) was produced and transduced using the same protocol described above and the cells were screened by fluorescence.

### LDH Release: Cell Viability

To measure the cell viability of the A549 single and double knockout cell lines, we performed a similar LDH assay as previously described (41). Approximately 2x10^4^ A549 wild-type, ΔRNF213, ΔRARRES3, or ΔRNF213/ΔRARRES3 cells were seeded into 96-well plates until ∼80% confluency. Afterward, the cells were stimulated with 20 U/mL of IFNγ or left unstimulated for 24h, followed by infection with the wild-type, knockout, or complemented parasites at MOI 2 and 3, utilizing a coupled plaque assay to ensure precise comparison among the parasite strains. Then, 4-6h post-infection (h.p.i.), 1 μM of Compound 1 (MBP146-78, MedChemExpress cat. no.188343-77-3, Monmouth Junction, NJ, USA) was added, and 24h post-infection, 50 μL of supernatant was used to measure LDH release using the Cytotoxicity Detection Kit (Roche, Basel, Switzerland) according to the manufacturer’s instructions. Absorbance (OD490) values were read after a 10-minute incubation in the kit reagents. The positive/lysis control was cells treated with 2% Triton X-100 in cell media solution and noted as 100 percent LDH release, while uninfected cells were noted as 0%. The percentage of LDH release for parasite-infected cells was calculated as: % LDH = (OD490 value of infected cells - OD490 value of uninfected cells)/(OD490 value of lysis control - OD490 value of uninfected cells), as in (41).

### Growth Assay: Parasites per Vacuole

A549 (wild-type, single knockouts, and double knockouts) cells were seeded into 12 mm glass coverslips in 24-well plates and grown to ∼80% confluency for 24h. The following day, they were left unstimulated or stimulated with 20-100 U/mL IFNγ for 24h. Wild-type, knockout/mutant, and complemented parasites were syringe-lysed, and 1x10^5^ parasites were used to infect each coverslip after IFNγ stimulation 1 h p.i. cells were washed three times with PBS to remove extracellular parasites, and they were incubated for an additional 23h. Afterward, the coverslips were fixed with 4% PFA for 20 minutes, permeabilized, and blocked in a buffer containing 3% (wt/vol) BSA, 5% (vol/vol) goat serum, and 0.1% Triton X-100 in PBS at room temperature for 1 h. Wild-type or mutant parasites that were not expressing GFP were stained with primary antibodies, anti-SAG1 (1:1000) (kindly provided by Dr. John C. Boothroyd, Stanford University) or anti-IMC1 (1:2000) (kindly provided by Dr. Gary Ward, University of Vermont), overnight at 4°C. The coverslips were then incubated with the following secondary antibodies for 1h at room temperature: anti-rabbit (1:3000) or anti-mouse (1:3000). Strains that were already GFP positive were not processed this way. Additionally, primary and secondary antibodies, rabbit anti-GRA7 (1:5000) and goat anti-rabbit 594 (Alexa Fluor, Thermo Fisher Scientific, cat. no.A-11007), were used as a marker for the PVM.

At least 100 vacuoles were counted for three biological replicates, and statistical analyses were performed using Python. Average parasites/vacuole values were analyzed using a three-way ANOVA (strain × treatment × cell type, Type II sums of squares). Simple effects (cell type within each strain/treatment, or treatment within each strain/cell type) were tested as linear contrasts from the fitted model, using its pooled residual error. P-values were corrected for multiple comparisons using the Benjamini-Hochberg (FDR) method, with significance set at α = 0.05, followed by graphical visualization in GraphPad Prism (Version 11.0.2 (100) for MacOS, GraphPad Software, Boston, Massachusetts USA, https://www.graphpad.com).

### Coating Assay: ISG Recruitment

A549 (wild-type, single knockouts, and double knockouts) cells were seeded into 12mm glass coverslips in 24-well plates and grown to ∼80% confluency for 24h. The following day, they were left unstimulated or stimulated with 20U/mL IFNγ for 24h. Wild-type, knockout/mutant, and complemented parasites were syringe-lysed, and 1x10^5^ parasites were used to infect each coverslip. 4 h.p.i., cells were washed three times with PBS to remove extracellular parasites, and they were reincubated for an additional 20h in the presence of Compound 1 for RARRES3 recruitment or fixed for ubiquitin and RNF213 staining. The coverslips were fixed with 4% PFA for 20 minutes, permeabilized, and blocked in a buffer containing 3% (wt/vol) BSA, 5% (vol/vol) goat serum, and 0.2% Triton X-100 in the blocking buffer (for anti-RNF213 and anti-ubiquitin) for 1h or 0.02% Saponin for 15 min (for anti-RARRES3) in PBS at room temperature. Only for A549 GFP and A549 overexpressing RARRES3 infected cells the fixation was made with ice-cold methanol for 10 min and permeabilization was also 0.02% Saponin for 15 min. Wild-type or mutant parasites that were not expressing GFP or if fixed with methanol were stained with primary antibodies, anti-SAG1 (1:1000) or anti-IMC1 (1:2000), overnight at 4°C. The coverslips were then incubated with the following secondary antibodies for 1h at room temperature: anti-rabbit 488 (Alexa Fluor, Thermo Fisher Scientific, cat. no.A-21206) (1:3000) or anti-mouse 488 (Alexa Fluor, Thermo Fisher Scientific, cat. no.A-21202) (1:3000) for 1h. Strains that were already GFP positive were not processed this way. Cells were also co-stained with primary antibodies, rabbit anti-RNF213 (1:100) (Sigma-Aldrich, cat. no.HPA026790), mouse anti-ubiquitin (1:250) (Enzo Life Sciences, Farmingdale, NY, USA, cat. no.ENZ-ABS840-0100), or rabbit anti-RARRES3 (1:1000) (Proteintech Group, Rosemont, IL, USA, cat. no.12065-1-AP). Additionally, primary and secondary antibodies, rabbit anti-GRA7 (1:5000), goat anti-rabbit 594 (Alexa Fluor, Thermo Fisher Scientific, cat. no.A-11007) or goat anti-rabbit 405 (Sigma-Aldrich, cat. no. SAB4600461), were used as a marker for the PVM. At least 100 vacuoles were counted for three biological replicates. Quantification of ISG recruitment was performed using a blinded method to prevent observer bias. Samples were randomly coded with numerical identifiers before microscope analysis, and the codes were revealed only after all data collection was completed. Vacuoles were considered positive when staining co-localized with the *Toxoplasma* PV, defined by a complete or partial ring around the vacuole. Vacuoles with no staining were categorized as negative. Fluorescence intensity was not considered in this analysis. Statistical analysis was performed using a 2-way ANOVA in GraphPad Prism.

### Western Blotting

Protein samples were prepared from either freshly extracellularly lysed tachyzoites or fully confluent cells. Parasites were harvested and centrifuged at 570 × g for 7 min at 4 °C. The parasite pellet or cells were lysed in ice-cold modified RIPA buffer (50 mM Tris-HCl pH 8.0, 150 mM NaCl, 1% NP-40, 0.5% sodium deoxycholate, and 0.1% SDS) supplemented with Halt™ Protease and Phosphatase Inhibitor Cocktail (Thermo Fisher Scientific, cat. no. 78442). Protein extracts equivalent to 1 × 10⁷ parasites or 1 × 10⁵ cells per lane were mixed in Laemmli sample buffer, denatured and resolved via 10% SDS-PAGE, transferred onto PVDF membranes, and blocked with PBS containing 5% non-fat dry milk. The membranes were incubated overnight with primary antibodies against HA (rat, 1:1000), SAG1 (rabbit, 1:5000), RNF213 (rabbit, 1:1000), or GAPDH (mouse, 1:1000; Santa Cruz Biotechnology, cat. no. sc-32233). After washing, membranes were probed with appropriate secondary antibodies (Thermo Fisher Scientific, 1:5000) and visualized using a ChemiDoc imaging system (BioRad).

### Bioinformatic Analysis

For GRA66 sequence analysis among strains the protein sequence of 15 *Toxoplasma gondii* strains were downloaded from ToxoDB database (42) (TGME49_320490, TGGT1_320490, TGVEG_320490, TGARI_320490, TGBR9_320490, TGCAST_320490, TGCOUG_320490, TGDOM2_320490, TGFOU_320490, TGMAS_320490, TGP89_320490, TGPRC2_320490, TGRH88_006720, TGRUB_320490, TGVAND_320490) and four orthologs *Besnoitia besnoiti* (BESB_043400), *Cystoisospora suis* (CSUI_008310), *Hammondia hammondi* (HHA_320490), *Sarcocystis neurona* (SRCN_3114) were submitted to alignment in PRALINE sequence alignment (43).

For prediction of signal peptide and transmembrane domains, sequence-based topology and signal peptides were analyzed using Phobius (44) under default parameters.

### Structural and docking analysis of GRA66 as a NAPE-PLD

The AlphaFold model of GRA66 (UniProt A0A125YX08) was retrieved from the AlphaFold Protein Structure Database (45) and trimmed to its folded metallo-beta-lactamase (MBL) domain (residues 350-800 for superposition; 430-785 for docking) on the basis of per-residue pLDDT and the positions of the MBL catalytic motifs. Global fold similarity to human NAPE-PLD (PDB 4QN9, chain A) (46) and to a panel of MBL-superfamily structures (glyoxalase II, 1QH5; CPSF73, 2I7T; NDM-1, 3SPU; tRNase Z, 1Y44) and an HKD-family phospholipase D (1F0I) was computed with TM-align (tmtools v0.3.0) (47), reporting TM-scores normalized by the reference length and Cα RMSD. Metal-coordinating residues of 4QN9 were defined as protein atoms within 2.8 Å of either catalytic Zn2+. Residue correspondence was established by global pairwise sequence alignment (BLOSUM62; Biopython v1.86) and confirmed structurally after a sequence-anchored, outlier-pruned superposition (Biotite v1.2); active-site conservation was quantified as the Cα RMSD over the seven mapped metal ligands. Lipid-channel residues were defined as protein atoms within 4.5 Å of the phosphatidylethanolamine (3PE) bound in 4QN9. Substrate binding was evaluated with Boltz-2 (48) by co-folding the GRA66 MBL domain (or the human NAPE-PLD sequence, or a His→Ala metal-site mutant) with two Zn^2+^ ions and either a truncated NAPE (SMILES CCCC(=O)OCC(OC(=O)CCC)COP(=O)(O)OCCNC(=O)CCCCC) or a truncated phosphatidylcholine control (SMILES CCCC(=O)OCC(COP(=O)([O-])OCC[N+](C)(C)C)OC(=O)CCC), using the automatic MSA server. Predictions were run both without restraints and with distance restraints between each Zn^2+^ and its mapped protein ligandsThe top-ranked model was analyzed for binuclear-zinc assembly (per-zinc protein coordination within 2.8 Å and Zn-Zn distance) and for engagement of the scissile phosphate (closest phosphate-oxygen-to-zinc distance). Predicted complexes were rendered, and molecular lipophilic potential surfaces computed, in UCSF ChimeraX v1.9 (49).

### Primers and sgRNA

Primers and sgRNAs used in this study are listed in table S2.

### *Mycoplasma* testing

Cell cultures were routinely tested for *Mycoplasma* contamination via PCR. Only *Mycoplasma*-free cultures were used in this study.

### Data availability statement

All data generated in this study are included in this manuscript and in its supplementary information file. Python scripts used in this study are available upon request.

## Results

### IFNγ susceptibility of Δ*gra66* and Δ*gra70* parasites is independent of RNF213

Previous CRISPR-Cas9 screens identified GRA66 and members of the GRA70 complex as parasite effectors required for optimal fitness in IFNγ-stimulated HFFs (16, 17). Loss of either GRA66 or GRA70 results in premature parasite egress and impaired intracellular growth, but the host restriction pathways responsible for these phenotypes remain unknown. Because the E3 ubiquitin ligase RNF213 is a major mediator of IFNγ-induced *Toxoplasma* restriction in human cells (28), we asked whether the increased susceptibility of *Δgra66* and *Δgra70* parasites is RNF213 dependent.

We first used CRISPR-Cas9 to reintroduce a C-terminally HA-tagged GRA66 into the *UPRT* locus of the *Δgra66* strain (16) (*Δgra66*::GRA66). To assess the requirement of GRA66’s predicted NAPE-PLD enzymatic activity, we also generated a catalytic-site mutant by substituting the first two histidines of the conserved HxHxDH zinc-binding motif with alanines (H540A/H542A) (*Δgra66*::GRA66mut). Expression of both tagged proteins was confirmed by immunoblotting (Fig. S1A), and the substitutions were verified by Sanger sequencing (Fig. S1C). GRA66 localizes to the PVM and PV lumen (16, 18) by immunofluorescence against the HA tag, *Δgra66*::GRA66mut and *Δgra66*::GRA66 showed the same distribution (Fig. S1B). For GRA70, we used previously characterized *Δgra70* and *Δgra70*::GRA70 strains (16, 17).

We first quantified parasite replication in A549 WT and *ΔRNF213* cells by scoring the number of parasites per vacuole in naive and IFNγ-stimulated cells (Fig. 1A). In A549 WT cells, IFNγ restricted the replication of all parasite strains, shifting the distribution toward smaller vacuoles (1-2 parasites). To compare the magnitude of restriction between the two cell lines directly, we expressed the IFNγ-induced loss of large vacuoles as the log2 fold decrease in the percentage of vacuoles containing eight or more parasites (Fig. 1B). In *ΔRNF213* cells, this restriction was significantly relieved for WT parasites and for the complemented strains *Δgra66*::GRA66 and *Δgra70*::GRA70, whereas it was unchanged for *Δgra66, Δgra66*::GRA66mut, and *Δgra70* parasites (Fig. 1B). Thus, WT parasites depend on RNF213 for the bulk of IFNγ-induced growth restriction, whereas the mutants remain restricted in its absence, and GRA66 requires its predicted catalytic activity to confer this protection.

**Figure 1.**
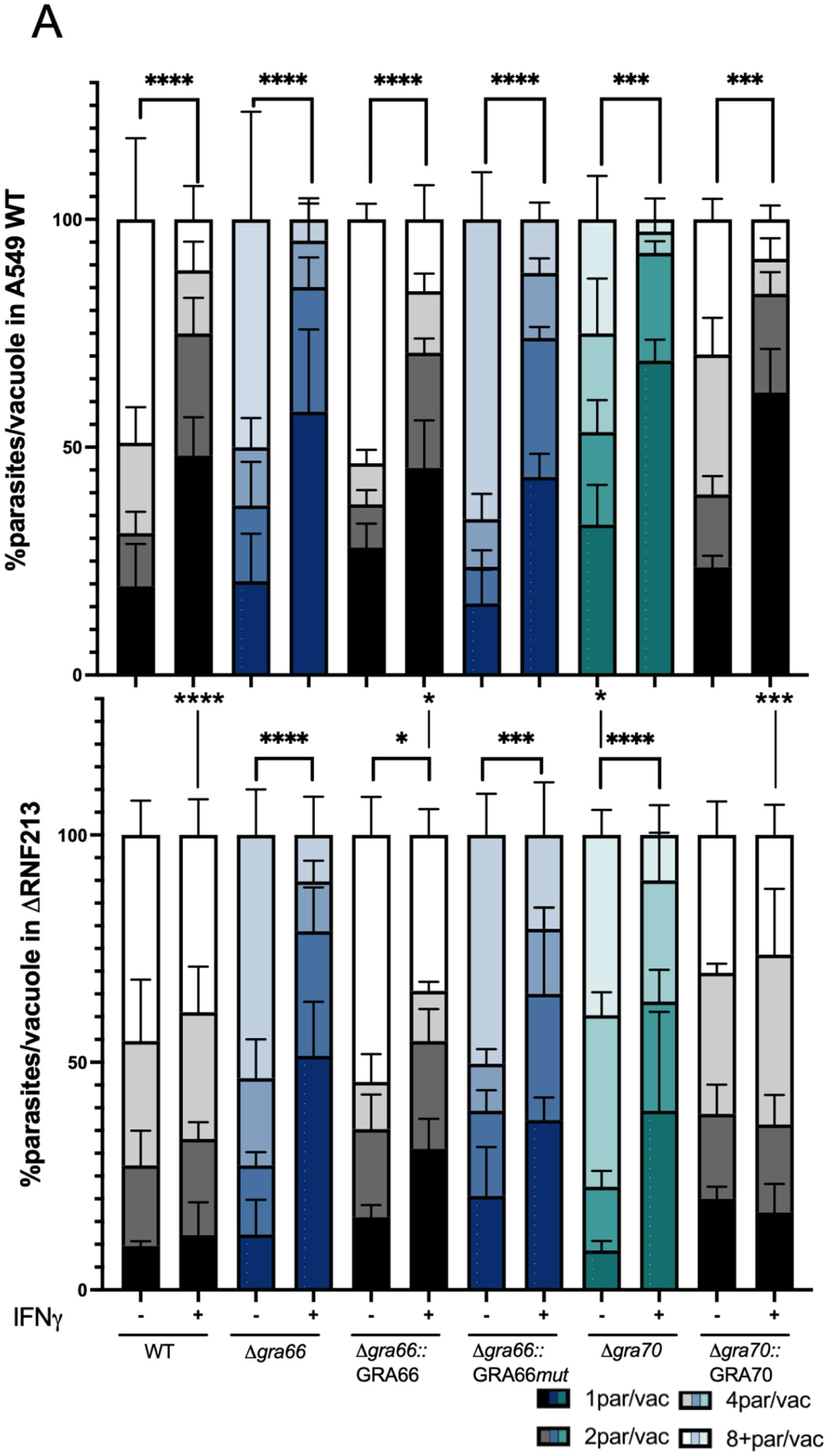
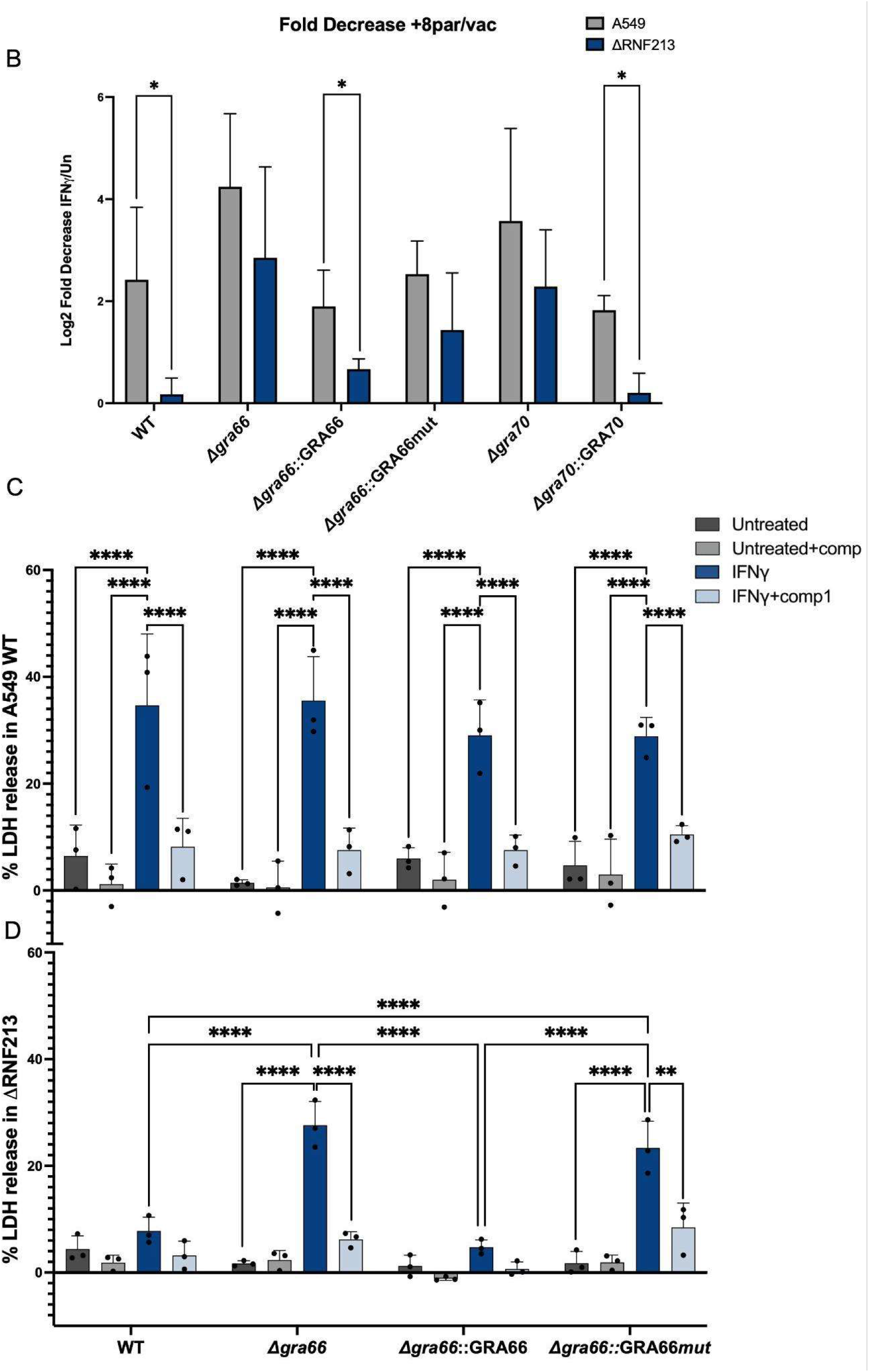
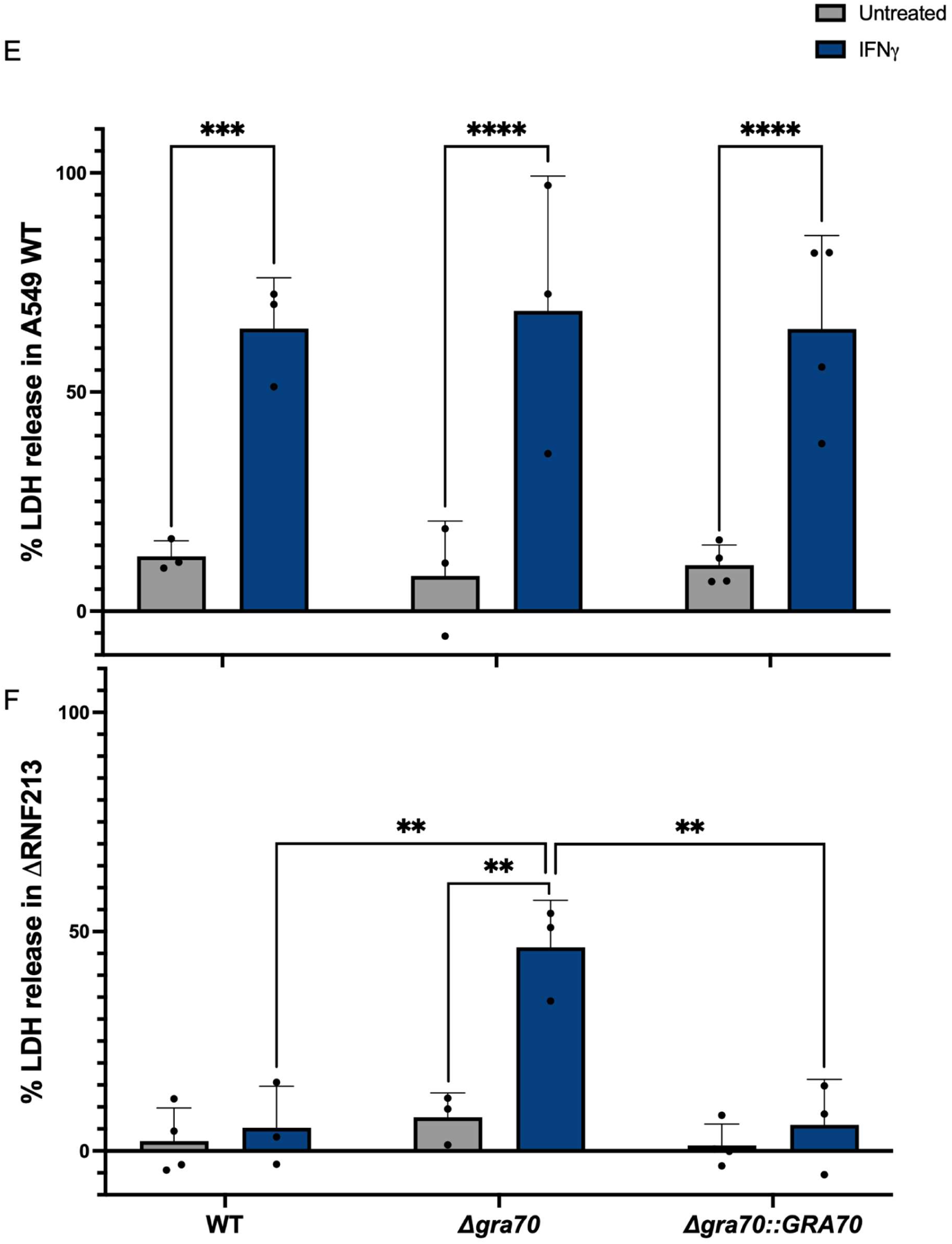
IFNγ susceptibility of *Δgra66* and *Δgra70* parasites is independent of RNF213. A549 WT and ΔRNF213 cells were left untreated or pre-stimulated with IFNγ (20 U/mL) for 24h prior to infection with the indicated *Toxoplasma* strains for 24h. (A) Intracellular parasite replication was assessed by quantifying the number of parasites per vacuole (≥100 vacuoles per condition). (B) IFNγ-induced loss of large vacuoles, expressed as the log2*-1 fold decrease in the percentage of vacuoles containing 8+ parasites (IFNγ/unstimulated); higher values indicate stronger restriction. (C-F) Host cell death quantified by LDH release assay at 24h post infection (h.p.i.) in A549 WT (C, E) and ΔRNF213 (D, F) cells infected with GRA66 knockout and complemented parasites (C, D) or GRA70 knockout and complemented parasites (E-F). To distinguish egress-driven lysis from primary host cell death, infections in (C) and (D) were treated with or without the PKG inhibitor Compound 1 (1µM) added at 6 h.p.i. to block active egress; this control was previously reported for the GRA70 knockout and complemented strains (16) and was therefore omitted from (E) and (F). Statistical significance for LDH release was determined by 2-way ANOVA followed by Tukey’s multiple comparison test. Significance levels for parasites per vacuole were calculated using the mean number of parasites per vacuole as a global continuous variable, assessed by 3-way ANOVA followed by the Benjamini-Hochberg (FDR) post-test (*P<0.05, **P<0.01, ***P<0.001, P****<0.0001. ns = non-significant).

Because impaired growth in these mutants is frequently linked to premature egress (16), we measured host cell death by lactate dehydrogenase (LDH) release, using the PKG inhibitor Compound 1 to block active parasite egress (50). In A549 WT cells, IFNγ stimulation induced significant LDH release across all tested strains, and this release was significantly reduced by Compound 1, indicating that it was driven by active parasite egress rather than by primary host cell lysis (Fig. 1C, E). In ΔRNF213 cells, IFNγ-induced LDH release upon WT infection was largely abrogated. In contrast, *Δgra66* and *Δgra66*::GRA66mut parasites (Fig. 1D), and *Δgra70* strains (Fig. 1F) still triggered significant egress-driven LDH release, which was reduced to WT levels in the complemented lines. Together, these results indicate that whereas RNF213 drives premature egress during WT infection, the IFNγ susceptibility of the *Δgra66* and *Δgra70* mutants is RNF213 independent and instead relies on a distinct restriction mechanism.

To further rule out RNF213 recruitment to the PV as the cause for *Δgra66* and *Δgra70* restriction, we quantified RNF213 and ubiquitin coating of the PVM. In IFNγ-stimulated A549 WT cells, RNF213 recruitment increased markedly on WT vacuoles but remained significantly lower on *Δgra66* and *Δgra66*::GRA66mut vacuoles, and complementation with wild-type GRA66 largely restored recruitment (Fig. 2A). A similar reduction in RNF213 recruitment was observed for *Δgra70* parasites in IFNγ-stimulated HFFs, the cell type in which this complex was originally characterized (Fig. 2C, Fig. S3), consistent with previous reports (17). Ubiquitin coating mirrored these RNF213 patterns: IFNγ markedly increased PVM ubiquitination in WT and complemented strains, whereas it remained low on *Δgra66*, *Δgra66*::GRA66mut, and *Δgra70* vacuoles (Fig. 2B, D), with a comparable trend already apparent in unstimulated cells. Notably, the catalytic-site mutant failed to restore either RNF213 or ubiquitin recruitment, indicating that the predicted enzymatic activity of GRA66 is required to generate or maintain the vacuolar features recognized by RNF213.

**Figure 2.**
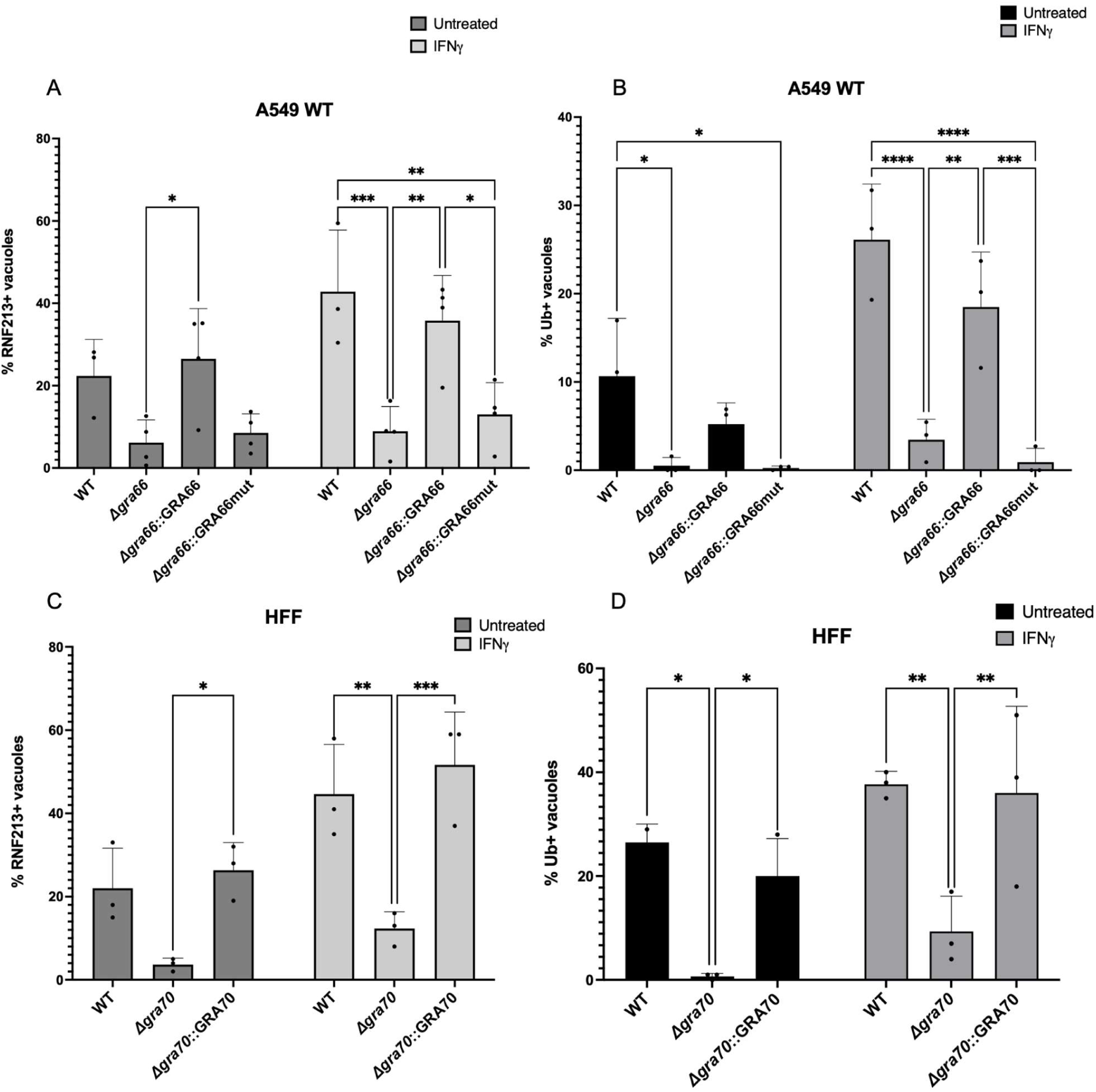
Vacuoles containing *Δgra66* or *Δgra70* parasites evade RNF213 recruitment and subsequent ubiquitination. A549 WT (A, B) or HFF (C, D) cells were left untreated or pre-stimulated with IFNγ (20U/mL) for 24h before infection with the indicated *Toxoplasma* strains. At 4 h.p.i., cells were fixed and immunostained for RNF213 (A, C) or ubiquitin (B, D). The percentage of positively coated vacuoles was determined by blindly counting at least 100 vacuoles per condition, scored as described in Material and Methods. Data represent the means ± SD of 3 independent experiments. Statistical significance was determined by 2-way ANOVA followed by Tukey’s multiple comparison test (*P<0.05, **P<0.01, ***P<0.001, P****<0.0001. ns = non-significant).

Together, these data show that Δ*gra66* and Δ*gra70* vacuoles are poorly recognized by RNF213 and are consequently poorly ubiquitinated. The increased IFNγ susceptibility of these mutants therefore cannot be explained by an exaggerated RNF213 response. Instead, loss of GRA66 renders parasites more susceptible to IFNγ while simultaneously reducing their recognition by RNF213, pointing to an alternative, RNF213-independent host restriction pathway.

### GRA66 prevents RARRES3-dependent premature egress

A recent overexpression screen identified the host phospholipase/acyltransferase RARRES3 as a restriction factor that drives premature *Toxoplasma* egress (31), a phenotype strikingly similar to what we observed for our *Δgra66* and *Δgra70* mutants upon IFNγ stimulation. Given that GRA66 is a putative NAPE-PLD, we hypothesized that GRA66 counteracts membrane changes produced by RARRES3 activity at the PVM.

To test whether endogenous RARRES3 is responsible for the susceptibility of Δ*gra66*, we assessed parasite-induced host cell death in CRISPR-generated ΔRARRES3 A549 cells. Deletion of RARRES3 rescued the premature egress phenotype of Δ*gra66* parasites: upon IFNγ stimulation, LDH release from Δ*gra66*-infected cells was significantly lower than from WT-infected cells and was restored by complementation with wild-type GRA66 but not with the catalytic mutant (Fig 3A). Because these cells retain an intact RNF213 pathway, WT and complemented parasites still elicited substantial LDH release, consistent with RNF213-driven egress in this background.

**Figure 3.**
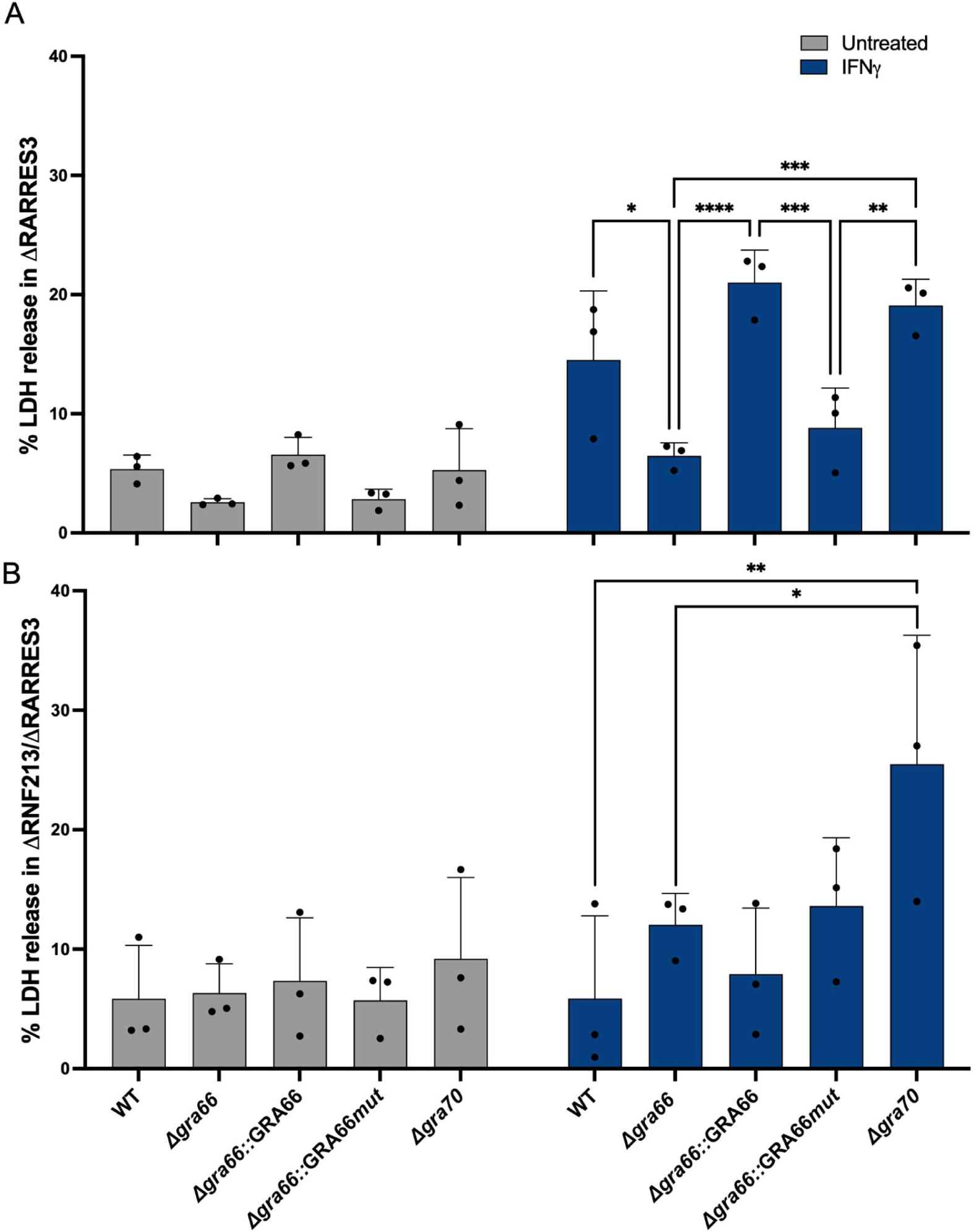
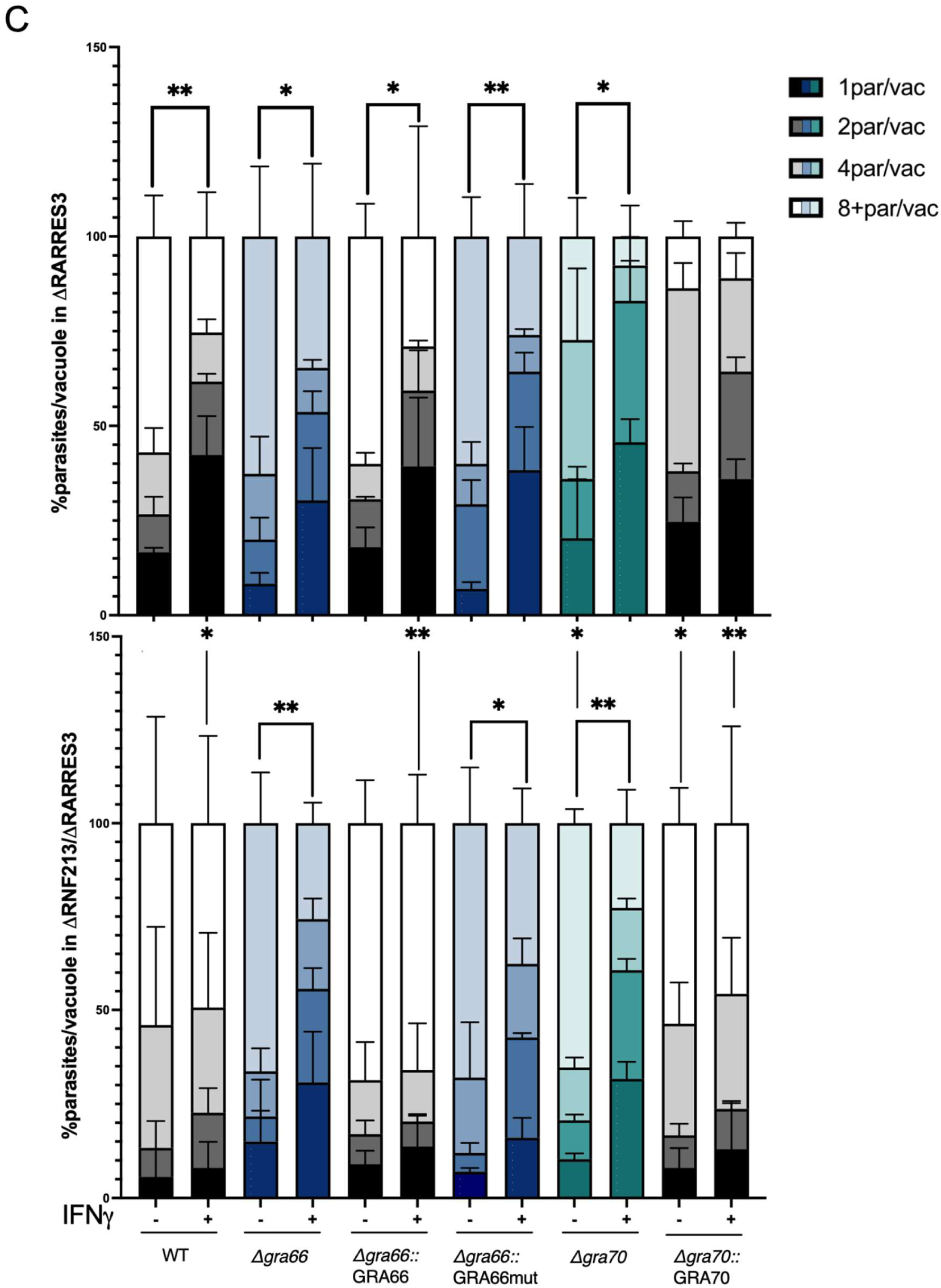
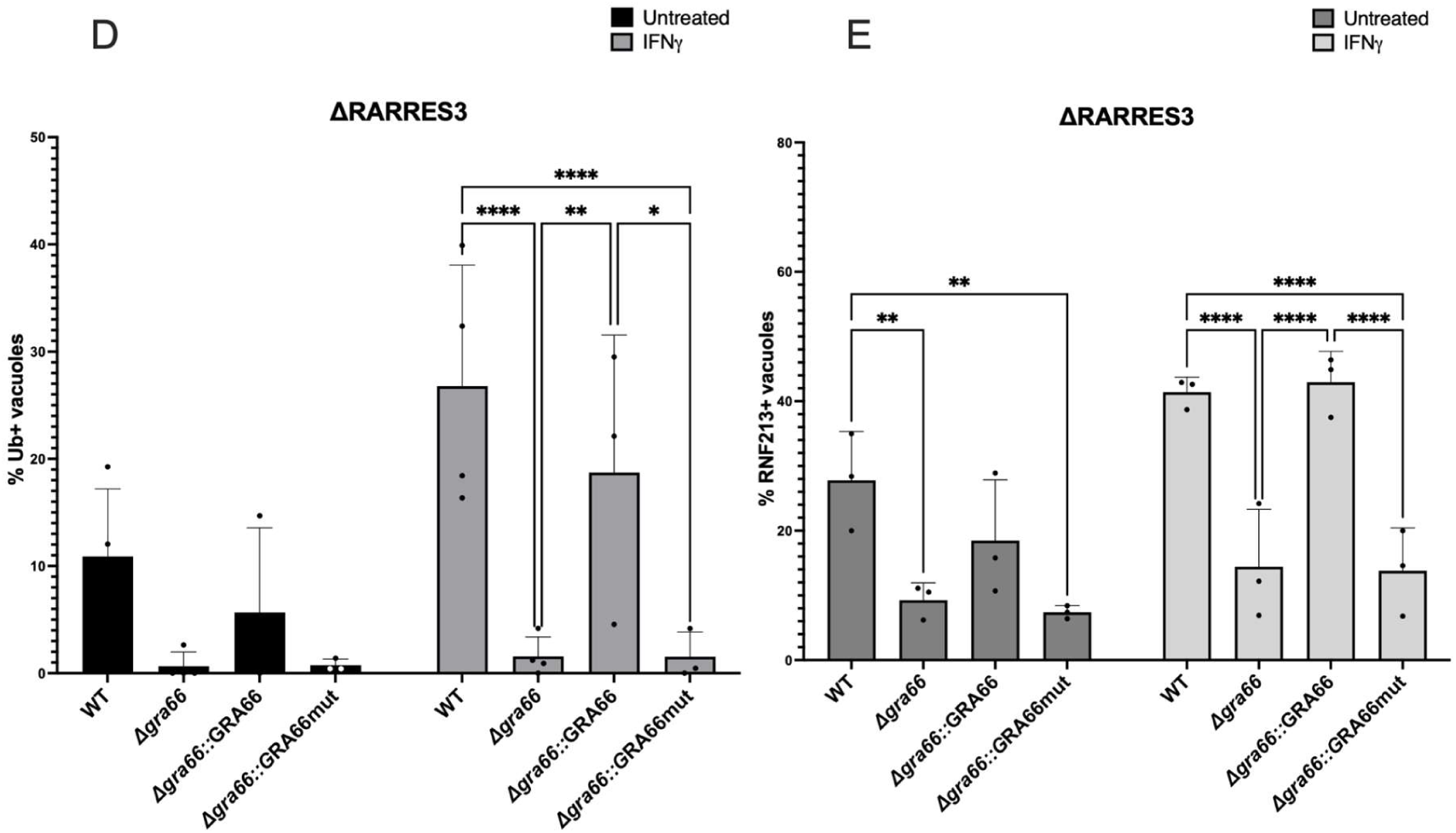
RARRES3 determines IFNγ-mediated egress of *Δgra66*, but not *Δgra70* parasites. (A, B) ΔRARRES3 (A) and ΔRNF213/ΔRARRES3 double-knockout (B) A549 cells were left untreated or pre-stimulated for 24h with IFNγ (20U/mL) prior to infection with the indicated *Toxoplasma* strains. LDH release was measured at 24h.p.i. to assess host cell death. (C) Intracellular parasite replication assessed by scoring the number of parasites per vacuole in unstimulated and IFNγ-stimulated cells. (D, E) ΔRARRES3 cells were pre-stimulated with IFNγ (20U/ml) for 24h, infected for 4 h, and scored for the percentage of vacuoles positive for ubiquitin (D) or RNF213 (E) by blindly counting of at least 100 vacuoles per condition, as described in Materials and Methods. Data represent the means ± SD of three independent experiments. Statistical significance was determined using a 2-way ANOVA followed by Tukey’s multiple comparison test (*P<0.05, **P<0.01. ns = non-significant). Significance levels for parasites per vacuole were calculated using the mean number of parasites per vacuole as a global continuous variable, assessed by 3-way ANOVA followed by the Benjamini-Hochberg (FDR) correction (*P<0.05, **P<0.01, ***P<0.001, P****<0.0001. ns = non-significant).

To uncouple this rescue from canonical restriction mechanisms, we generated ΔRNF213/ΔRARRES3 double-knockout cells (Fig S2). In these cells, IFNγ-induced LDH release from *Δgra66*- and *Δgra66*::GRA66mut-infected cells was no longer significantly different from WT infection (Fig. 3B). In contrast, cell death induced by the *Δgra70* mutant was unaffected by loss of RARRES3 in either the single or double knockout lines and remained significantly above WT (Fig. 3B). This divergence separates the two RNF213-independent restriction pathways and places GRA66, but not the GRA70 complex, in the same pathway as RARRES3 (Fig. 3B).

While RARRES3 deletion prevented premature egress of *Δgra66* parasites, we asked whether it also restored intracellular parasite replication. By quantifying parasites per vacuole, we found that all parasite strains still exhibited reduced growth in IFNγ-stimulated single ΔRARRES3 cells. In the ΔRNF213/ΔRARRES3 double knockouts, replication of WT and complemented strains was largely restored, whereas *Δgra66* and *Δgra66*::GRA66mut parasites remained significantly restricted (Fig. 3C). This residual defect is consistent with the mislocalization of GRA23, a pore-forming protein implicated in host nutrient acquisition (Fig. S4), although we have not directly tested whether reduced nutrient acquisition accounts for it (51). Thus, while *Δgra66* parasites undergo RARRES3-driven egress, loss of GRA66 also causes a secondary, egress-independent and RARRES3-independent growth defect whose mechanism remains untested and may involve additional IFNγ-induced restriction pathways.

To confirm that RARRES3 does not affect RNF213-mediated restriction, we quantified RNF213 and ubiquitin recruitment to the PVM in ΔRARRES3 cells. As in WT A549 cells, ΔRARRES3 cells infected with Δ*gra66* parasites displayed significantly reduced PVM coating with RNF213 and ubiquitin; this defect was rescued by complementation with wild-type GRA66 but not by the catalytic-site mutant (Fig. 3D, E). The lower LDH release during Δ*gra66* infection of ΔRARRES3 cells is therefore not explained by altered RNF213 recruitment.

To assess the impact of RARRES3 levels on parasite survival, we used a lentiviral system to transduce A549 WT and ΔRARRES3 cells, subsequently isolating single clones stably expressing either a GFP control or RARRES3. Because A549 WT cells retain an intact RNF213 response, WT parasites elicited high levels of host cell death even in the GFP control lines (Fig. 4A). In contrast, Δ*gra66* parasites, which evade RNF213 recruitment (Fig 2), caused little host cell death in ΔRARRES3 + GFP clones (Fig. 4B). Restoring RARRES3 expression in these cells significantly increased host cell death during *Δgra66* infection, re-establishing the egress phenotype (Fig. 4B); the catalytic-site mutant behaved identically (Fig. 4D), confirming that the predicted enzymatic activity of GRA66 is required to counter RARRES3. We next examined RARRES3 overexpression in the A549 WT clones. Whereas WT parasites resist baseline endogenous RARRES3, lentiviral overexpression of the host phospholipase significantly increased host cell death, and the same trend was observed for *Δgra66::*GRA66 (Fig. 4A, C). Conversely, RARRES3 overexpression did not significantly exacerbate death of *Δgra66*-infected cells relative to baseline WT A549 cells (Fig. 4B), likely because these mutants are already maximally susceptible to endogenous RARRES3 levels. Together, these data support a model in which GRA66 is required to protect against RARRES3-dependent premature egress, operating in parallel to the canonical RNF213 pathway.

**Figure 4.**
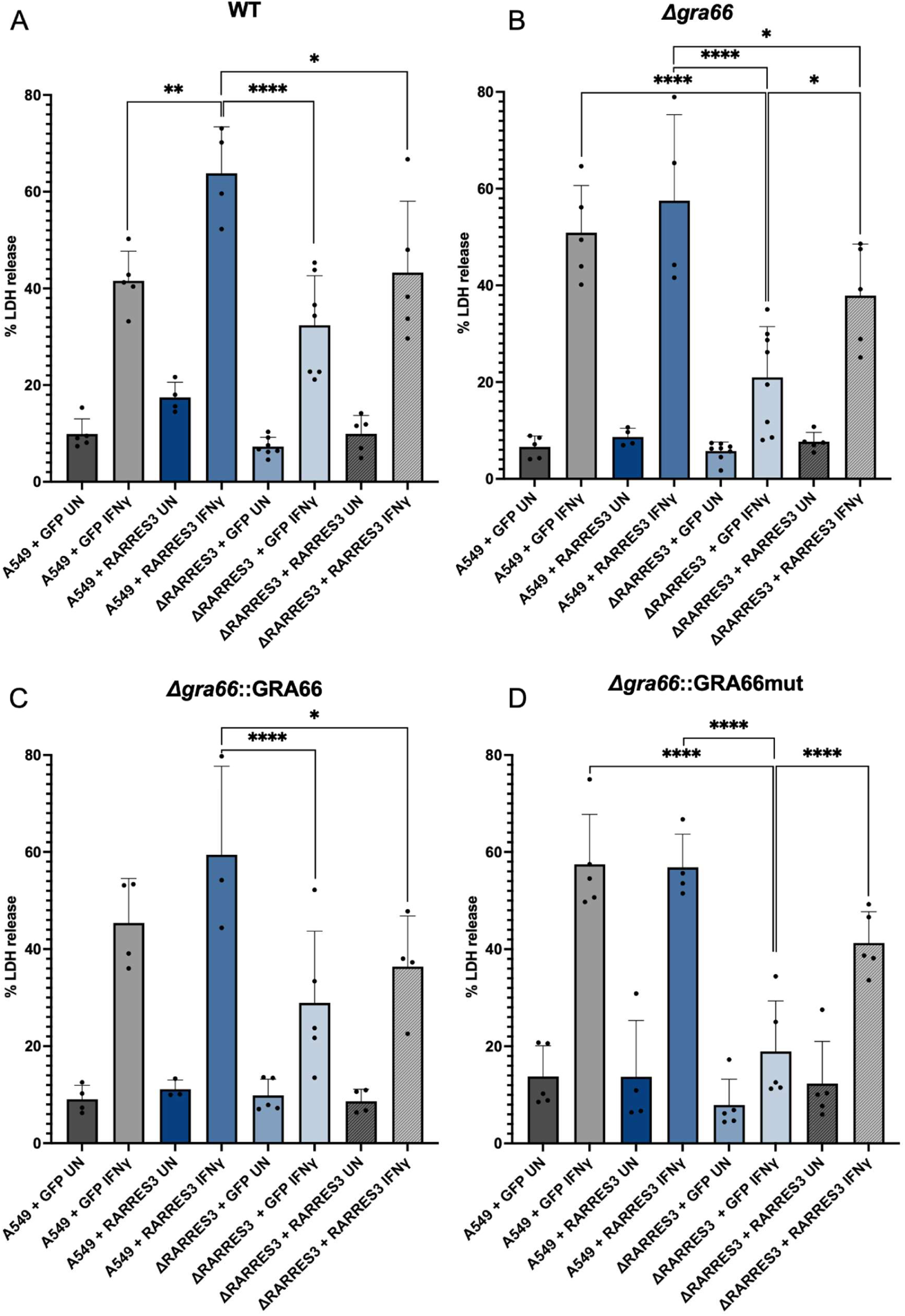
RARRES3 expression levels determine host cell death during *Δgra66* infection. Single clones of A549 WT and ΔRARRES3 cells stably expressing either a GFP control or RARRES3 following lentiviral transduction were left untreated (UN) or pre-stimulated with IFNγ (20U/mL) for 24h. Cells were then infected with WT (A), *Δgra66* (B), *Δgra66*::GRA66 (C), or *Δgra66*::GRA66mut (D) parasites and host cell death was quantified by LDH release at 24h.p.i. Data represent the means ± SD of at least three independent experiments. Statistical analyses were performed using 2-way ANOVA, followed by Tukey’s multiple comparison test (*P<0.05, **P<0.01, ***P<0.001, P****<0.0001. ns = non-significant).

### RARRES3 is recruited to the *Toxoplasma* PV following IFNγ stimulation

To determine whether RARRES3 acts at the PVM, we assessed its subcellular localization during infection. By immunofluorescence microscopy of IFNγ-stimulated cells, RARRES3 was recruited to the PV and the intravacuolar network (IVN), where it colocalized with the *Toxoplasma* dense granule protein GRA7 (Fig. 5A). These results place RARRES3 at the vacuolar membranes, where its phospholipase and acyltransferase activities could act on PVM lipids.

**Figure 5.**
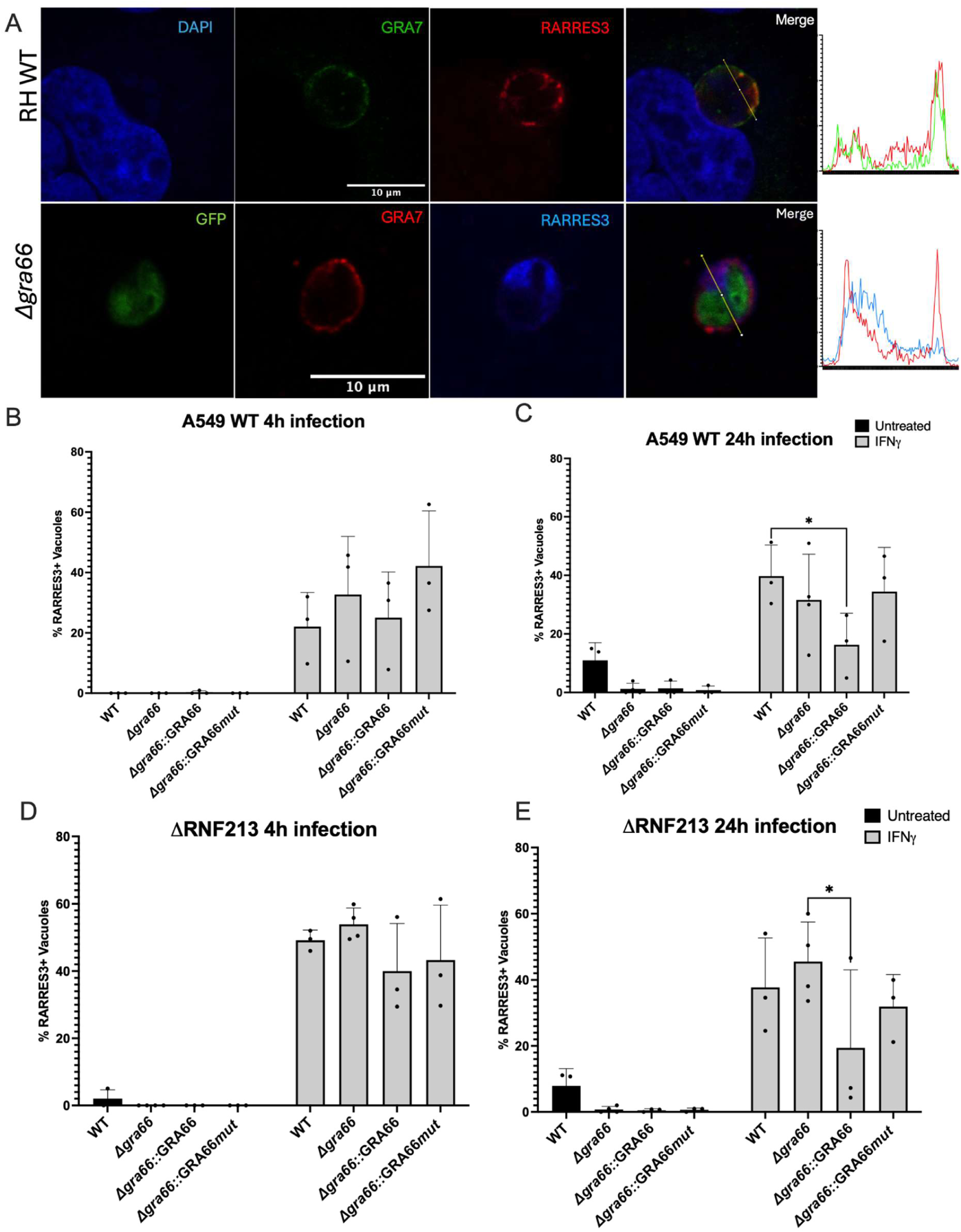
RARRES3 targets the PV independently of RNF213. (A) A549 WT cells were pre-stimulated with IFNγ (20 U/ml) for 24h and infected with *Toxoplasma* WT or *Δgra66* parasites. At 24 h.p.i., cells were fixed and immunostained for RARRES3 and the PVM/IVN marker GRA7. Note that the fluorophore assignments differ between rows because *Δgra66* parasites constitutively express GFP: in the RH WT row GRA7 is shown in green and RARRES3 in red, whereas in the *Δgra66* row GFP is green, GRA7 is red and RARRES3 is blue. Line profiles at right show fluorescence intensity for the two indicated channels along the arrow drawn in the merged image. Scale bar, 10 μm. (B-E) A549 WT (B, C) or ΔRNF213 (D, E) cells were pre-stimulated with IFNγ (20 U/ml) for 24h, infected with the indicated parasite strains and assessed for RARRES3 recruitment to the PV at 4 h.p.i. (B, D) and 24h.p.i. (C, E). For the 24h infections, Compound 1 (1 μM) was added 4 h.p.i to prevent premature egress. The percentage of RARRES3-coated vacuoles was determined by blindly counting of at least 100 vacuoles per condition, scored as described in Material and Methods. Data represent the means ± SD of three independent experiments. Statistical significance was determined by 2-way ANOVA followed by Tukey’s multiple comparison test (*P<0.05, **P<0.01, ***P<0.001, P****<0.0001. ns = non-significant). Scale bar, 10 μm.

We next quantified RARRES3 recruitment across parasite strains. In cells expressing endogenous RARRES3, recruitment was negligible in the absence of IFNγ and was strongly induced by IFNγ on vacuoles of all strains, in both A549 WT and *ΔRNF213* cells and at both 4 and 24 h.p.i. (Fig. 5B-E), indicating that recruitment under these conditions does not require GRA66. RARRES3 recruitment was slightly higher in *ΔRNF213* (Fig. 5D). At 24 h.p.i., RARRES3 recruitment was lower on *Δgra66*::GRA66 vacuoles, reaching significance in A549 WT (Fig. 5C) and *ΔRNF213* cells (Fig. 5E). RARRES3 therefore reaches the vacuole irrespective of GRA66, and functional GRA66 does not prevent, and may modestly reduce, its accumulation.

Having established that IFNγ drives RARRES3 to the PV, likely because it is not expressed without IFNγ, we asked whether elevating RARRES3 levels alone was sufficient to trigger recruitment. Using A549 clones overexpressing RARRES3, we quantified PV coating at 24h.p.i. in the absence or presence of IFNγ stimulation, adding compound 1 to prevent egress. Unexpectedly, in unstimulated cells, overexpression produced substantial RARRES3 recruitment only during infection with WT or GRA66-complemented parasites, whereas vacuoles containing containing *Δgra66* or the catalytic mutant remained largely devoid of RARRES3 (Fig. 6A, B). This GRA66 dependence was specific to unstimulated conditions: upon IFNγ stimulation, RARRES3 was recruited to vacuoles of all four strains, with no reduction on *Δgra66* or the catalytic-mutant vacuoles (Fig. 6B).

**Figure 6.**
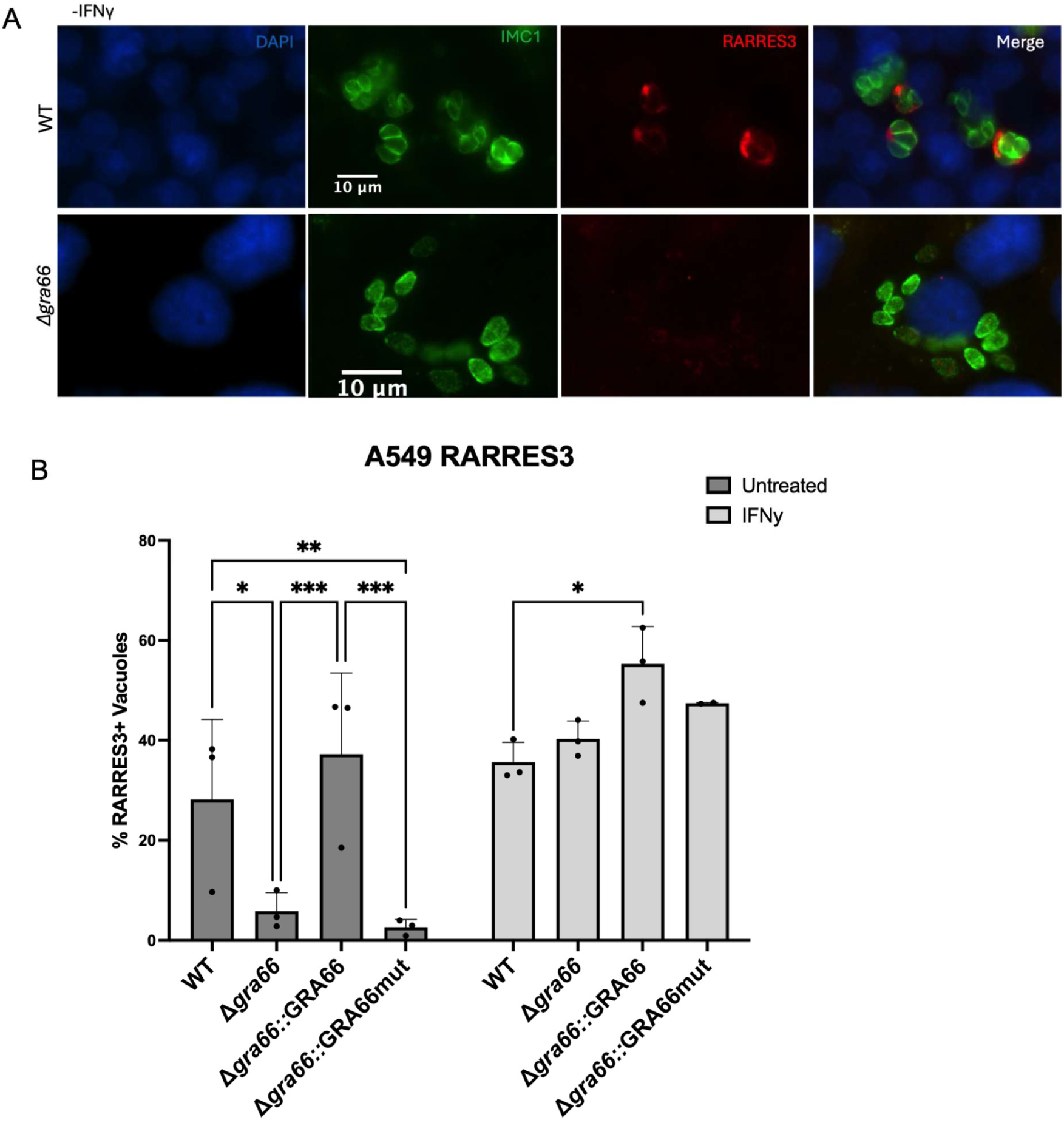
RARRES3 recruitment in unstimulated cells required functional GRA66. (A) Representative images of A549 cells overexpressing RARRES3, left unstimulated, infected with WT or *Δgra66* parasites for 24h and immunostained for RARRES3 and the parasite marker IMC1; DAPI marks nuclei. Images from IFNγ stimulated cells and from the complemented and catalytic-mutant strains are quantified in (B). Scale bar, 10 μm. (B) Quantification of RARRES3 recruitment to the PV at 24h.p.i. in unstimulated and IFNγ-stimulated (20 U/ml, 24h) A549 RARRES3-overexpressing cells infected with the indicated strains. Compound 1 (1 μM) was added at 4 h.p.i to block active egress. The percentage of RARRES3-coated vacuoles was determined by blindly counting of at least 100 vacuoles per condition. Data represent the means ± SD of three independent experiments. Statistical significance was determined by 2-way ANOVA followed by Tukey’s multiple comparison test (*P<0.05, **P<0.01, ***P<0.001, P****<0.0001. ns = non-significant).

These two experiments therefore reveal opposite dependencies. Under IFNγ stimulation, RARRES3 recruitment occurs independently of GRA66 and is, if anything, reduced by functional GRA66; in unstimulated cells with elevated RARRES3, recruitment requires functional GRA66. Together these observations suggest that RARRES3 reaches the PVM by more than one route, only one of which depends on a membrane feature generated or maintained by GRA66 (see Discussion).

### *In silico* structural analysis supports GRA66’s predicted NAPE-PLD activity

Given that GRA66 is a putative NAPE-PLD, we asked whether its predicted structure supports this assignment. GRA66 is a 1,134-residue protein that is mostly predicted to be disordered but contains a single well-folded domain (residues ∼430–785) with a classic metallo-β-lactamase (MBL) fold, including the diagnostic HxHxDH zinc-binding motif (H540-N-H542-P-D544-H545). Superimposing this domain onto the crystal structure of human NAPE-PLD (PDB 4QN9) (46) gave a TM-score of 0.88 and a Cα RMSD of 2.0 Å over 314 aligned residues sharing 41.7% sequence identity, values indicating homology rather than a coincidental family resemblance. Comparison with other MBL-superfamily enzymes of unrelated activity showed that GRA66 resembles NAPE-PLD (TM 0.88) more closely than tRNase Z (0.59), NDM-1 β-lactamase (0.56), glyoxalase II (0.52), or CPSF73 (0.47), and falls well below the TM=0.5 threshold for a shared fold when compared with the mechanistically distinct HKD-family phospholipases D (0.28; Fig. 7A). GRA66 therefore belongs to the NAPE-PLD-type phospholipase D family of its proposed host counterpart rather than to the more commonly studied HKD-PLD family.

**Figure 7.**
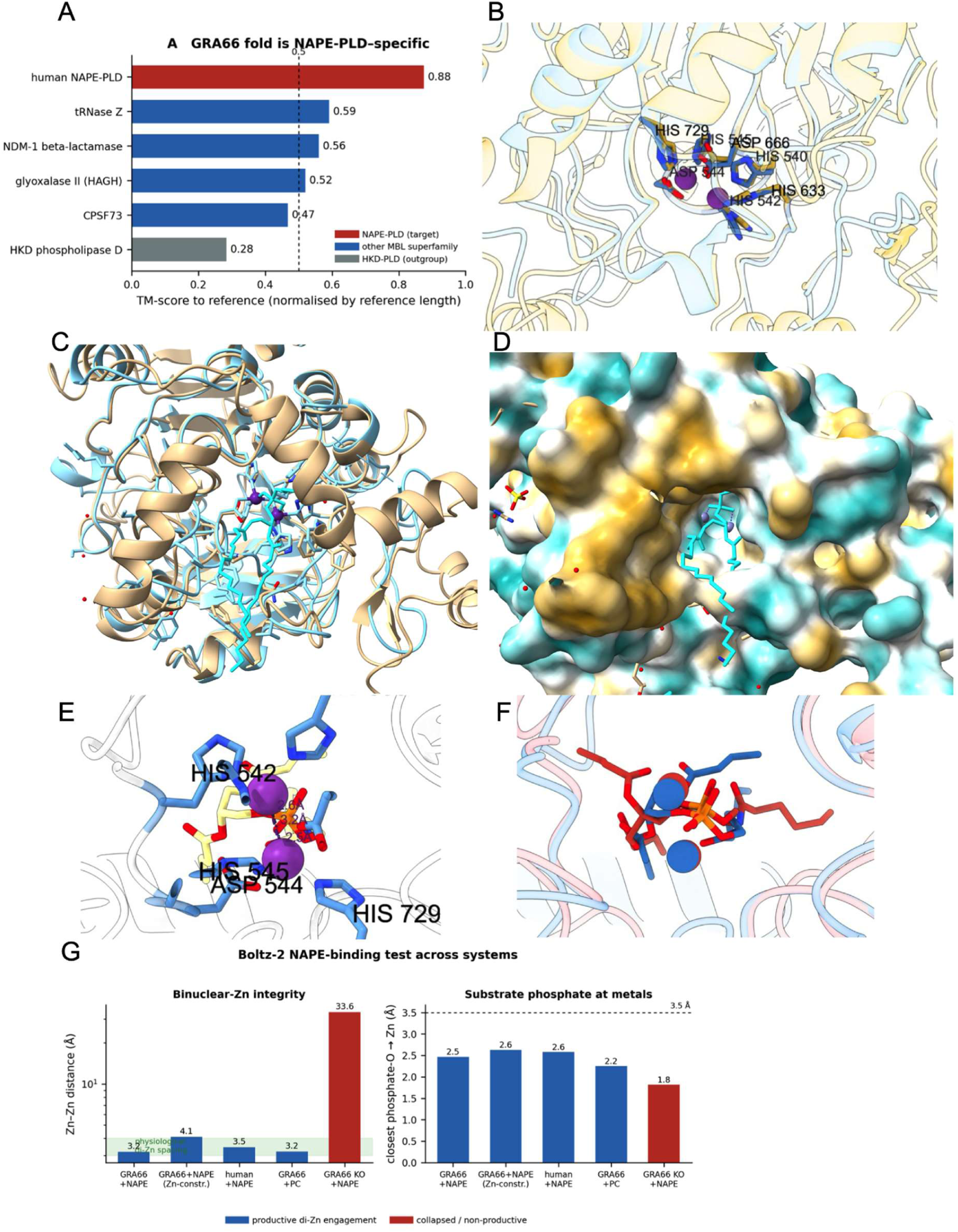
GRA66 shares a conserved catalytic architecture with human NAPE-PLD. All structures shown are computational models or model-to-crystal superpositions. (A) TM-scores of the GRA66 MBL domain superimposed on human NAPE-PLD (red; PDB 4QN9) and on representative MBL-superfamily enzymes (blue) and an unrelated HKD-family phospholipase D (grey); dashed line, TM = 0.5. (B) Superposition of the GRA66 AlphaFold model (khaki) on human NAPE-PLD (blue) showing the conserved di-zinc active site (GRA66 numbering shown); crystallographic Zn²⁺ ions frp, 4QN9 are shown as purple spheres. (C) Superposition of the GRA66 (khaki) model onto the human NAPE-PLD crystal structure (blue) showing convergence on the same substrate pocket, occupied in 4QN9 by a bound phosphatidylethanolamine (3PE, cyan sticks); catalytic Zn²⁺ ions shown as purple spheres. (D) Molecular lipophilic potential surface of GRA66 (hydrophobic, gold; hydrophilic, teal). The 3PE acyl chain (cyan sticks) is the ligand from the 4QN9 crystal structure, positioned on the GRA66 surface by the superposition in (C), and illustrates the location of the conserved hydrophobic channel. (E) Top-ranked Boltz-2 model of the GRA66 MBL domain (grey cartoon) co-folded with a NAPE ligand (khaki carbons, red oxygens, orange phosphorus) and two Zn²⁺ ions (purple); interface pTM = 0.95. (F) Overlay of the GRA66 (blue) and human NAPE-PLD (red) Boltz-2 complexes after protein superposition, showing coincidence of the docked NAPE and di-zinc centers. (G) Zn-Zn distance (left) and closest phosphate-oxygen-to-zinc distance (right) across docking systems: GRA66+NAPE (unrestrained and Zn-restrained), the human-enzyme positive control, GRA66+phosphatidylcholine, and the GRA66 metal-site knockout. Blue, productive di-zinc engagement; red, collapsed/non-productive. Note that for the metal-site knockout the phosphate-oxygen distance is measured to the single remaining zinc ion, which is displaced from the active site (Zn-Zn 33.6 Å); the short value therefore does not indicate productive engagement.

The predicted active site supports this assignment. All seven metal-coordinating residues of human NAPE-PLD are conserved in GRA66 and superimpose closely (Cα RMSD 0.27 Å): His185, His187, His253, and Asp284, which together coordinate the first zinc ion, correspond to His540, His542, His633 and Asp666 in GRA66, while Asp189, His190 and His343, which coordinate the second zinc, correspond to Asp544, His545 and His729 (Fig. 7B). GRA66 thus retains an intact, correctly arranged di-zinc site, in contrast to MBL-fold proteins that have lost metal-binding residues and are catalytically inactive.

The two proteins also converge beyond the active site. Overlaying the GRA66 model onto the human NAPE-PLD crystal structure shows that both fold around the same pocket, occupied in 4QN9 by a bound phosphatidylethanolamine (PE) molecule (Fig. 7C), indicating a shared substrate-binding architecture rather than only a catalytic center. Consistent with this, the hydrophobic channel that accommodates the phospholipid acyl chain is conserved at the sequence level: of the 21 residues lining the PE in the human structure, 76% are identical or conservatively hydrophobic in GRA66, including Trp634 and Arg637. Mapping the molecular lipophilic potential onto the GRA66 surface shows this channel as a continuous hydrophobic groove running alongside the acyl tail of the superimposed lipid (Fig. 7D), matching the shape expected for an acyl-chain-binding phospholipase. Notably, the 28 strain-polymorphic residues we identified across *Toxoplasma* lineages (Fig. S6A) all fall outside this active site and channel, consistent with both being under strong functional constraint.

To get insights into whether GRA66 could potentially engage its proposed substrate, we used Boltz-2 (48) to co-fold the GRA66 MBL domain together with two Zn²⁺ ions and a NAPE molecule. GRA66 assembled a canonical binuclear-zinc center (Zn-Zn distance 3.2 Å) and positioned the substrate so that a phosphate oxygen bridged both metals (2.6 and 2.5 Å; Fig. 7E), with high model confidence (interface pTM 0.95). Equivalent predictions in which the zinc ions were restrained to their mapped protein ligands gave the same arrangement (Fig. 7G), indicating that the result does not depend on unguided metal placement. This geometry closely mirrored that obtained for human NAPE-PLD under the same conditions (Zn-Zn 3.4 Å; phosphate oxygenes 2.8 and 2.6 Å), and overlaying the two predicted complexes showed that the docked NAPE and both zinc ions aligned within 1 Å of their human counterparts (Fig. 7F). The arrangement required an intact metal site: substituting the five zinc-coordinating histidines (H540A/H542A/H545A/H633A/H729A, a more extensive substitution than the H540A/H542A parasite mutant, chosen to ablate both metal sites) collapsed the binuclear center, with the Zn-Zn distance increasing to 33.6 Å and loss of any bridging interaction (Fig. 7G). In control predictions, phosphatidylcholine also reached the di-zinc center through its shared glycerophosphate backbone (Fig. 7G), indicating that this assay reports substrate engagement and catalytic-site competence rather than headgroup selectivity. Together, these structural analyses support the assignment of GRA66 as a catalytically competent NAPE-PLD, consistent with the requirement for its predicted enzymatic activity that we observed throughout this study.

### Working model for GRA66-RARRES3 antagonism at the PVM

Our results suggest a model (Fig. 8) in which RARRES3 is recruited to the PVM upon IFNγ stimulation independently of GRA66. At the PVM, RARRES3 remodels vacuolar phospholipids through its Ca^2+^-independent N-acetyltransferase activity, transferring an acyl chain from a donor phospholipid to the primary amine of phosphatidylethanolamine (PE) to generate N-acylphosphatidylethanolamine (NAPE) and releasing a single-tailed lysophospholipid, lysophosphatidylcholine or lysophosphatidylethanolamine (LPC/LPE), as a coupled reaction product (32, 33). Both products are predicted to alter membrane biophysics. LPC and LPE, which bear a single acyl tail and a comparatively large polar head, favor positive membrane curvature and behave as mild detergents, properties associated with pore formation and membrane fission (52, 53). NAPE, in contrast, carries an additional acyl chain relative to its polar head and is predicted to favor negative curvature, following the shape-based principles established for cone-shaped membrane lipids (54). Because both species impose a spontaneous curvature that differs from that of the surrounding bilayer, local accumulation of either is expected to introduce curvature stress and destabilize the PVM.

**Figure 8.**
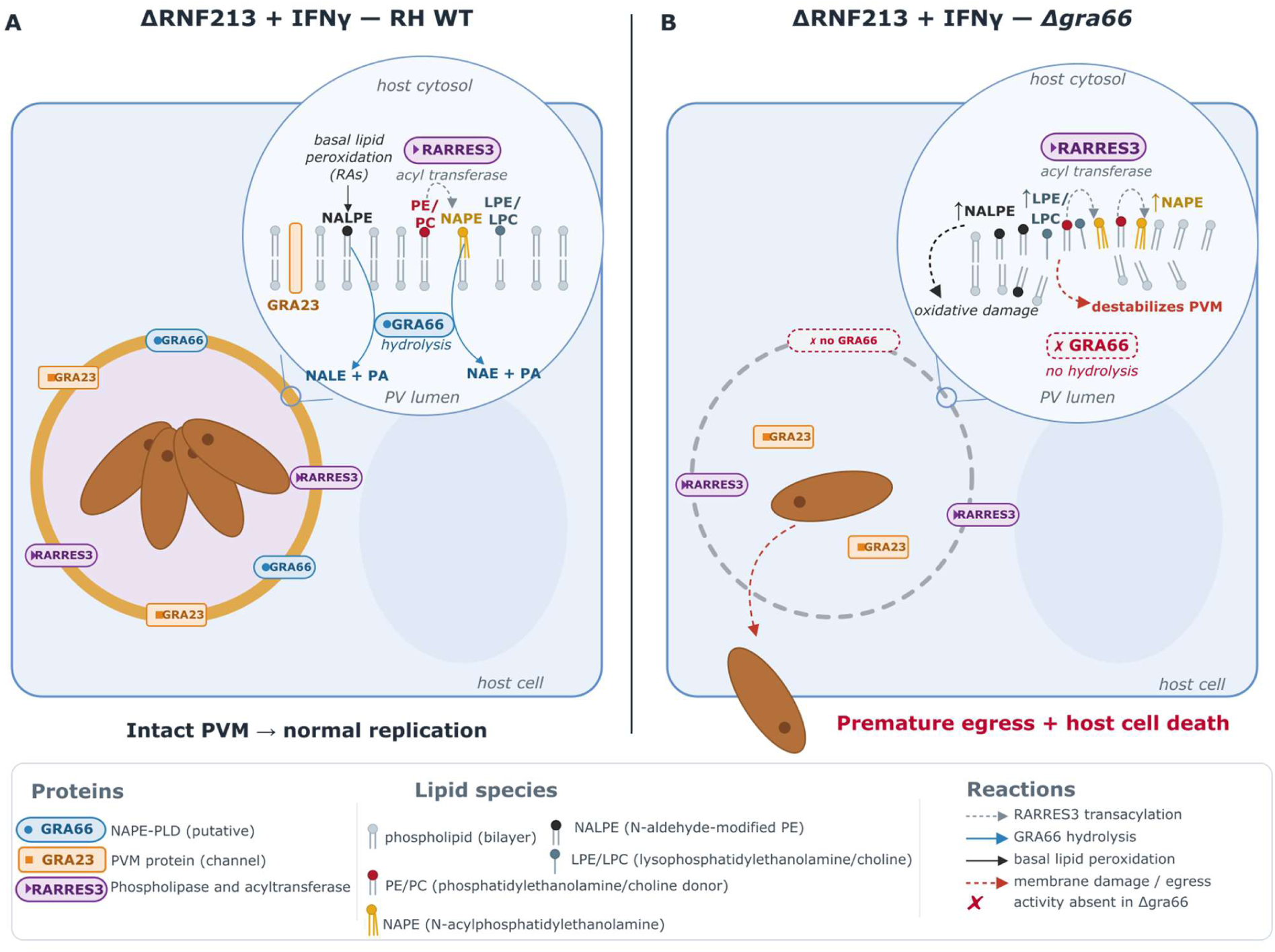
Proposed model for GRA66-dependent protection of the PVM, independent of RNF213. The model is depicted in ΔRNF213 A549 cells stimulated with IFNγ, the background in which the RARRES3-dependent pathway can be observed in isolation. (A) In WT parasites, RARRES3 is recruited to the PVM, where its N-acyltransferase activity transfers an acyl chain from a donor phospholipid to the primary amine of phosphatidylethanolamine (PE), generating NAPE and releasing a lysophospholipid (LPC/LPE). GRA66, a predicted NAPE-PLD, hydrolyzes NAPE to N-acylethanolamine (NAE) and phosphatidic acid (PA), and may additionally hydrolyze N-aldehyde-modified PEs (NALPEs) arising from basal lipid peroxidation. This activity is proposed to preserve PVM integrity, supporting normal intracellular parasite replication and correct insertion of PVM proteins. (B) In *Δgra66* parasites, RARRES3 is still recruited to the PVM and continues to generate NAPE and LPC/LPE, but NAPE is no longer cleared; LPC/LPE is not a NAPE-PLD substrate and persists in both conditions. Accumulation of NAPE at the PVM is proposed to destabilize the membrane, resulting in premature parasite egress and host cell death through a mechanism independent of RNF213-mediated ubiquitination.

We propose that in WT parasites GRA66 hydrolyzes the NAPE generated by RARRES3, limiting its accumulation and preserving the vacuolar niche required for parasite replication and survival (Fig. 8A). Mammalian NAPE-PLD also efficiently hydrolyzes N-aldehyde modified-PEs (NALPEs) generated by basal lipid peroxidation (55), raising the possibility that GRA66 additionally clears such adducts from the PVM, although we have not tested this. We note that the products of NAPE hydrolysis, N-acetylethanolamine and phosphatidic acid, are themselves bioactive lipids, so GRA66 may shape PVM lipid signaling as well as limit substrate accumulation.

In the absence of GRA66, we hypothesize that unhydrolyzed NAPE accumulates at the PVM and compromises its integrity, providing a plausible trigger for the premature egress we observe (Fig. 8B). The LPC/LPE generated by the phospholipase activity of RARRES3 would not be a substrate for a NAPE-PLD and would therefore persist irrespective of GRA66. We have not directly measured PVM permeability or integrity in this study, and this step of the model, whether RARRES3 activity destabilizes the PVM in the absence of GRA66, remains to be tested directly, for example by assessing PVM permeability to a small tracer in the presence of Compound 1 to uncouple membrane integrity from egress itself.

## Discussion

Our data identify the *Toxoplasm*a dense granule protein GRA66 as a parasite effector required to prevent premature egress driven by the host phospholipase and acyltransferase RARRES3, independently of RNF213. GRA66 was dispensable for parasite replication in unstimulated cells but was required for fitness once cells were exposed to IFNγ, indicating that it is not a general growth factor but is specifically required to withstand an IFNγ-induced challenge. Although GRA66 and the GRA70 complex were both recovered as parasite factors required for fitness in IFNγ-stimulated human cells (16, 17), the two act through separable mechanisms: deletion of RARRES3 rescued the premature egress of *Δgra66* and of the catalytic-motif mutant, but not of *Δgra70*. GRA66 therefore counteracts a discrete, RARRES3-dependent process, and the shared premature egress phenotype of the two mutant classes reflects a convergence on a common outcome rather than a common host target.

We favor a model in which GRA66 and RARRES3 act on overlapping lipid substrates at the PVM, in opposing directions. RARRES3 belongs to the PLAAT family, whose members possess both phospholipase A1/A2 activity, which generates lysophospholipids, and a Ca^2+^-independent N-acyltransferase activity, which transfers an acyl chain from a donor phospholipid to the primary amine of phosphatidylethanolamine to generate N-acylphosphatidylethanolamine (NAPE) (32, 33). NAPE is the substrate of the NAPE-hydrolyzing phospholipase D family, to which GRA66 is predicted to belong on the basis of its metallo-beta-lactamase fold, its zinc-binding HxHxDH motif, and its structural correspondence to human NAPE-PLD (PDB 4QN9; Fig. 7). This suggests a concrete and testable point of biochemical opposition: RARRES3 delivered to the PVM after IFNγ stimulation remodels vacuolar glycerophospholipids and, through its N-acyltransferase activity, generates NAPE, whereas GRA66 hydrolyzes NAPE and thereby limits accumulation of this product at the vacuolar membrane. Such a scheme accounts for observations that are otherwise unconnected: that catalysis is required on both sides, since a catalytically dead RARRES3 does not restrict (31) and our catalytic-motif GRA66 mutant does not complement; that the requirement is specific to RARRES3 and not to the GRA70 complex; and that GRA66 acts downstream of RARRES3 recruitment rather than preventing it.

We present this as a working hypothesis rather than a demonstrated pathway, and note three limitations. First, the relative contribution of N-acyltransferase and phospholipase activities of RARRES3 at the PVM is unknown; if the restrictive species is instead a lysophospholipid generated by the phospholipase activity, GRA66 would not directly reverse it. Second, neither enzyme has been characterized biochemically in this system, so the predicted activity of GRA66 remains inferred from sequence and structure. Third, premature egress is itself a parasite-driven process, and at the level of an individual vacuole it may represent an escape response to an increasingly inhospitable niche rather than a host-driven killing mechanism analogous to IRG/GBP-mediated vacuole lysis.

The relationship between RARRES3 recruitment and RARRES3 function requires explanation, because our recruitment experiments give apparently opposite results. After IFNγ stimulation, RARRES3 was recruited to vacuoles of all strains and, if anything, accumulated slightly less on vacuoles containing functional (Fig. 5B, E). By contrast, in unstimulated cells overexpressing RARRES3, recruitment occurred on wild-type and complemented vacuoles but was largely absent from Δ*gra66* and catalytic-mutant vacuoles (Fig. 6). Furthermore, Δ*gra66* vacuoles are compositionally abnormal even without immune stimulation, as shown by mislocalization of the transmembrane pore GRA23 (Fig. S4). The simplest reconciliation is that RARRES3 reaches the PVM by more than one route: a dominant, GRA66-independent route engaged by IFNγ, which may saturate the response, and a second, lower-capacity route that depends on a membrane feature which generated or maintained by functional GRA66 and that becomes rate-limiting only when IFNγ is absent.

It was shown that GRA66 is a substrate of ASP5 and in the absence of cleavage it is not located properly in the PVM, also lack of ASP5 mislocalizes several GRAs, a phenotype we can not exclude happens through GRA66 (18, 56). An alternative interpretation deserves equal weight. PLAAT3, a related family member, is recruited to organelle membranes in response to small-pore membrane damage (57), raising the possibility that RARRES3 recruitment reports pre-existing membrane perturbation rather than initiating it. Our genetic data establish that RARRES3 is required for the premature egress of Δ*gra66* parasites and is therefore causal for the phenotype, but they do not distinguish whether RARRES3 initiates PVM damage or amplifies damage that arises from another IFNγ-induced insult. Resolving initiation from amplification, for example by testing whether a catalytically inactive RARRES3 is still recruited, will be important for placing RARRES3 correctly within the pathway.

This model raises the question of the substrate source for GRA66 in the absence of RARRES3. RARRES3 is an IFNγ-inducible enzyme, but it is not the sole source of modified PE in the cell: other PLAAT family members provide constitutive Ca^2+^-independent N-Acyltransferase activity, and Ca^2+^-dependent N-acyltransferases represent a further potential source, so basal substrates could be scavenged during PVM biogenesis. In addition, mammalian NAPE-PLD efficiently hydrolyzes N-aldehyde-modified PEs (NALPEs), toxic adducts generated during basal lipid peroxidation (55). The basal function of GRA66 may therefore extend to continuous membrane maintenance, clearing oxidatively damaged or aberrantly acylated host lipids to preserve a PVM able to support integral proteins such as GRA23. Upon IFNγ stimulation, upregulated RARRES3 may generate NAPE, or initiate parallel phospholipase-dependent remodeling, at a rate that exceeds the clearance capacity of GRA66.

A further observation is that Δ*gra66* and Δ*gra70* vacuoles recruited less RNF213 and were less ubiquitinated than wild-type vacuoles, yet were more restricted overall. Loss of GRA66 therefore allows the parasite to evade the RNF213 pathway while rendering it more susceptible to RARRES3, indicating that the two host pathways are not redundant and may read different features of the vacuole. RNF213 has a demonstrated capacity to act on lipid substrates, since it ubiquitylates the lipid A moiety of lipopolysaccharide on cytosolic *Salmonella* (58); it also coats phylogenetically diverse pathogens through a reaction that requires ATP hydrolysis by its dynein-like domain (27). If the ligand that RNF213 recognizes on the *Toxoplasma* PVM is a lipid or a lipid-dependent surface, a parasite lipid-modifying enzyme could plausibly influence its abundance, which would reconcile the reduced recruitment with a role for GRA66 in shaping PVM lipid composition. A simpler explanation is that loss of GRA66 disrupts insertion of PVM proteins in general, as indicated by the mislocalization of GRA23, and thereby alters whatever surface RNF213 detects, without any direct action on the RNF213 ligand. Either way, the reduced RNF213 recruitment argues against the idea that Δ*gra66* susceptibility reflects an exaggerated RNF213 response and supports the conclusion that the mechanism described here is RNF213 independent.

The persistence of a growth defect in Δ*gra66* parasites even in ΔRNF213/ΔRARRES3 cells, despite rescue of premature egress, indicates that GRA66 has functions beyond countering RARRES3. The mislocalization of GRA23 from the PVM to the vacuolar lumen in Δ*gra66* parasites (Fig. S4) offers a likely basis. GRA23, together with GRA17, forms non-selective pores that permit diffusion of small molecules across the PVM (12), and additional dense granule proteins including GRA47 and GRA72 influence the same permeability (11). Besides influencing nutrient acquisition and permeability, GRA23 and GRA72 were also found to impact parasite fitness under IFNγ stimulation (16). Impaired positioning of GRA23 would be expected to reduce the parasite’s access to host-derived small molecules, a deficit that would be aggravated under IFNγ stimulation, when human cells restrict availability of nutrients such as tryptophan through induction of indoleamine 2,3-dioxygenase (59). This nutritional hypothesis is consistent with the known permeability functions of GRA23 but has not been tested directly in our system; we cannot currently exclude that loss of GRA66 renders parasites susceptible to a distinct, unidentified IFNγ-induced restriction pathway unrelated to nutrient acquisition.

The strain distribution of RARRES3 sensitivity remains to be defined precisely. Our study establishes that RARRES3 restricts the type I RH strain, extending the pathway beyond the type III strain context in which RARRES3 was originally described (31). Whether susceptibility differs quantitatively among the clonal lineages is a separate question that our data do not resolve. If the type III lineage proves more sensitive, our sequence analysis argues against the simplest explanation: the nineteen polymorphic residues that distinguish the type III (VEG) GRA66 allele from the type I and type II alleles all lie outside the predicted catalytic site (Fig. S6), so a catalytically weaker type III enzyme is unlikely to be the cause. Differences in expression, stability, or localization of the GRA66 allele would be more plausible candidates and would be worth testing directly. Polymorphic parasite effectors are known to set the threshold of survival against host restriction factors, as shown for ROP5 and ROP18 in the murine system (60), and GRA66 may function analogously for RARRES3, although we note that the ROP5 and ROP18 example concerns murine rather than human immunity.

Our findings place lipid remodeling among the arms of human cell-autonomous immunity against *Toxoplasma*. IFNγ is increasingly recognized to reshape host lipid metabolism, for example through induction of cholesterol 25-hydroxylase (CH25H) and production of the antimicrobial oxysterol 25-hydroxycholesterol (25HC), although CH25H induction is driven predominantly by type I interferon (61, 62). Phospholipid remodeling represents a distinct axis (54, 63). RARRES3 generates lysophospholipids such as lysophosphatidylcholine (LPC), which alters membrane curvature and stability in model membranes, and it is plausible, though not established here, that localized lysophospholipid production can contribute to the loss of PVM integrity that we propose precedes premature egress. Consistent with a membrane-perturbing activity, LPC promotes P2X7 receptor-mediated responses associated with sustained Ca²⁺ influx and pore formation (52, 53). That RARRES3 overexpression produces only minimal changes in the global cellular lipidome during viral infection (35) is consistent with a mechanism based on localized rather than bulk membrane remodeling.

Similar dependencies on host phospholipids have been described in *Plasmodium*, where host-derived LPC regulates parasite metabolism and developmental progression (64) and host PC is required for correct localization of vacuolar membrane proteins during liver stage infection (65). RARRES3 has additionally been reported to modulate host signaling pathways, including PI3K/Akt, Wnt and β-catenin (66, 67), and mTOR (35), and we cannot exclude a contribution from such signaling to the restriction (68) we observe; however, the requirement for RARRES3 catalytic activity and its localization to the PVM lead us to favor a direct membrane-remodeling mechanism.

Collectively, our results support a model in which human non-immune cells deploy the lipid-remodeling enzyme RARRES3 at the *Toxoplasma* vacuole, where it is proposed to compromise vacuolar integrity and force premature parasite egress. The parasite counters this activity with GRA66, a predicted NAPE-hydrolizing phospholipase D that preserves the vacuolar niche. This defines a lipid-centered layer of cell-autonomous immunity that operates in parallel with, and independently of, the RNF213-mediated ubiquitination pathway, and it identifies the parasite effector required to withstand it. Direct measurement of PVM integrity, biochemical characterization of both enzymes, and lipidomic analysis of the vacuolar membrane will be needed to test the substrate relationship we propose and to establish whether the two enzymes act, as we suspect, on opposite sides of the same lipid.

## Acknowledgements

We thank Dr. David Sibley (Washington University in St. Louis) for providing the A549 ΔRARRES3 cells, the sgRNA plasmid used to generate this line, the lentiviral packaging plasmids (pMD2G, pRSV-Rev, and pMDLg/RRE), and the TRIP.RARRES3, TRIP.GFP and LentiCrispr_RARRES3 plasmids. We thank Dr. John C. Boothroyd (Stanford University) for the anti-GRA7 and anti-SAG1 antibodies, and Dr. Gary Ward (University of Vermont) for the anti-IMC1 antibody. We acknowledge Dr Bennet Penn (University of California Davis) for providing the Lenti-X 293T cells and Dr. Tatsunori Masatani (Gifu University, Japan) for anti-GRA23 antibodies. We also thank Ingrid Brust-Mascher and the UC Davis School of Veterinary Medicine Advanced Imaging Facility for confocal microscopy support and the staff at VEuPathDB for maintaining the database that made this work possible.

## Author contributions

J.P.J.S. and J.C. were responsible for project conceptualization, experiment design, and funding. E.A.J. made the initial observation of lack of host cell death in ΔRNF213 cells. C.M.S. performed experiments on figure 1 (A, C and D), figure 2 (A and B), figure 3, Figure 4, 5, 6 and 7 (C and D), and supplemental figures (S1, S2, S3, S4, S5 and S6). S.M.B. performed experiments for Figure 1 (A, E and F), figure 2 (C and D), figure 3 (C) and Figure S3. J.P.J.S. generated figure 7 (A, B, E, F and G). J.P.J.S, C.M.S., and S.M.B. wrote the paper and contributed to all analyses. J.P.J.S and J. C. revised the manuscript.

